# Exploring the distribution of phylogenetic networks generated under a birth-death-hybridization process

**DOI:** 10.1101/2022.11.10.516033

**Authors:** Joshua A. Justison, Tracy A. Heath

## Abstract

Gene-flow processes such as hybridization and introgression play important roles in shaping diversity across the tree of life. Recent studies extending birth-death models have made it possible to investigate patterns of reticulation in a macroevolutionary context. These models allow for different macroevolutionary patterns of gene flow events that can either add, maintain, or remove lineages—with the gene flow itself possibly being dependent on the relatedness between species—thus creating complex diversification scenarios. Further, many reticulate phylogenetic inference methods assume specific reticulation structures or phylogenies belonging to certain network classes. However, the distributions of phylogenetic networks under reticulate birth-death processes are poorly characterized, and it is unknown whether they violate common methodological assumptions. We use simulation techniques to explore phylogenetic network space under a birth-death-hybridization process where the hybridization rate can have a linear dependence on genetic distance. Specifically, we measured the number of lineages through time and role of hybridization in diversification along with the proportion of phylogenetic networks that belong to commonly used network classes (e.g., tree-child, tree-based, or level-1 networks). We find that the growth of phylogenetic networks and class membership are largely affected by assumptions about macroevolutionary patterns of gene flow. In accordance with previous studies, a lower proportion of networks belonged to these classes based on type and density of reticulate events. However, under a birth-death-hybridization process, these factors form an antagonistic relationship; the type of reticulation events that cause high membership proportions also lead to the highest reticulation density, consequently lowering the overall proportion of phylogenies in some classes. Further, we observed that genetic distance–dependent gene flow and incomplete sampling increase the proportion of class membership, primarily due to having fewer reticulate events. Our results can inform studies if their biological expectations of gene flow are associated with evolutionary histories that satisfy the assumptions of current methodology and aid in finding phylogenetic classes that are relevant for methods development.

## 1 Introduction

Gene flow—the exchange of genetic material between species—has been observed in a number of systems (Taylor and Larson 2019) and plays an important role in shaping Earth’s biodiversity (Stebbins 1959; Bock 2010; Mallet et al. 2016). Phylogenetic networks are used to directly infer reticulate histories and are vital in informing our understanding of adaptive radiations (Ottenburghs et al. 2016), mimicry patterns (Edelman et al. 2019), and even the phylodynamics of infectious diseases like COVID-19 (Wang et al. 2022). The field has recently seen an explosion of new methods for estimating phylogenetic networks (Elworth et al. 2019). However, due to the complex topological landscape of phylogenetic networks, robust model-based techniques remain limited in their ability to scale to large datasets (Hejase and Liu 2016). The birth-death-hybridization process (Morin and Moret 2006; Woodhams et al. 2016; Zhang et al. 2018) is an extension of a model commonly used to describe lineage diversification (*i.e*., the birth-death process, Kendall 1948; Nee et al. 1994b) and provides a statistical framework for understanding the generation of phylogenetic networks. With this model, we can understand how reticulate processes influence patterns of diversification and affect the distribution of species across the tree of life.

Birth-death processes are used to model the phylogenetic branching process on a macroevolutionary scale, thus revealing expected patterns of speciation and extinction (Kendall 1948; Raup 1985; Nee et al. 1994b; Nee 2006). A rich theoretical framework for this process allows us to describe the distribution of branch lengths (Steel and Mooers 2010; Mooers et al. 2012), growth dynamics (Nee et al. 1994a; Stadler 2008), and topological properties (Mooers and Heard 1997; Lambert and Stadler 2013) of phylogenetic trees. The birth-death family of models has proven useful for estimating diversification rates (Nee et al. 1994b,a; Höhna et al. 2015) and choosing biologically motivated priors in Bayesian analyses (Rannala and Yang 1996; Velasco 2008). However, analyses of divergence times (Pybus and Harvey 2000) and tree shape (Mooers and Heard 1997; Heath et al. 2008b; Jones 2011) have shown that empirical phylogenies differ from those expected under a constant rate birth-death process. To more accurately explain the observed patterns, these statistical models have undergone several extensions to account for a wide range of biological processes and systematic sampling challenges.

The birth-death-hybridization process (formally described below) generates phylogenetic networks under a process that allows lineages to undergo speciation, extinction, and reticulation (hybridization). Although— for eponymous consistency—events are denoted as hybridizations, the process does not make any mechanistic assumptions about gene flow; reticulate events on phylogenetic networks can model the many processes where genetic material is exchanged between lineages. Consequently, phylogenetic network inference has been used to describe histories of introgression (Thawornwattana et al. 2018; Myers et al. 2022), hybridization (Morales-Briones et al. 2018; Dolinay et al. 2021), and lateral gene transfer (Betat et al. 2015). The specific biological interpretation of a given reticulation event is a challenge that depends largely on the system under investigation and is further complicated by how reticulation events are modeled on the phylogenetic network (Hibbins and Hahn 2022).

Even though phylogenetic network methods are largely agnostic to the underlying cause of reticulation, existing implementations of the birth-death-hybridization process make different assumptions about how reticulation events are represented on the network. We denote each type of reticulation—lineage generative, lineage neutral, and lineage degenerative—based on how the events change the number of lineages on the phylogenetic network (Box 1). The structure and timing of each reticulation type can provide some biological meaning to reticulation events (Box 1; but see Hibbins and Hahn 2022). Consequently, caution is needed when trying to infer and interpret reticulate patterns; the lineage effects from the types of reticulations can have far reaching implications on species diversification dynamics.

**Box 1: Description of Reticulation Types**

Patterns of gene flow can take different shapes on phylogenetic networks. We differentiate types of reticulation depending on changes in the number of lineages on the phylogeny after an event (Fig. 1).

- **Lineage Generative**: A new lineage is created (denoted as m-type in Janssen and Liu 2021)
- **Lineage Neutral**: The number of lineages remains constant (n-type)
- **Lineage Degenerative**: The process has one fewer lineages after the event (y-type)

The table below outlines possible interpretations for each reticulate pattern. For example, lineage generative events are consistent with hybrid speciation or allopolyploidization since these events directly create new hybrid lineages. Introgression or lateral gene transfer can be seen as lineage neutral events as genetic material is exchanged, but the number of lineages remains constant. Interestingly, the formation of a new hybrid lineage also has potential to be lineage neutral or even lineage degenerative if genetic swamping occurs (Todesco et al. 2016). Genetic swamping is the loss of parental lineages after hybridization, resulting from either the hybrid having a higher fitness or a breakdown of reproductive barriers which eventually merges the hybrid lineage with its parental lineages. Extinction and incomplete sampling can further complicate the interpretation of reticulation events. Failure to sample the donor lineages of reticulation—so called ghost lineages (Ottenburghs 2020; Tricou et al. 2022)—may change its structure on the reconstructed phylogenetic network (Fig. S1). Without complete sampling, lineage generative and lineage neutral hybridization can appear as a different types of reticulation events.

**Figure 1:**
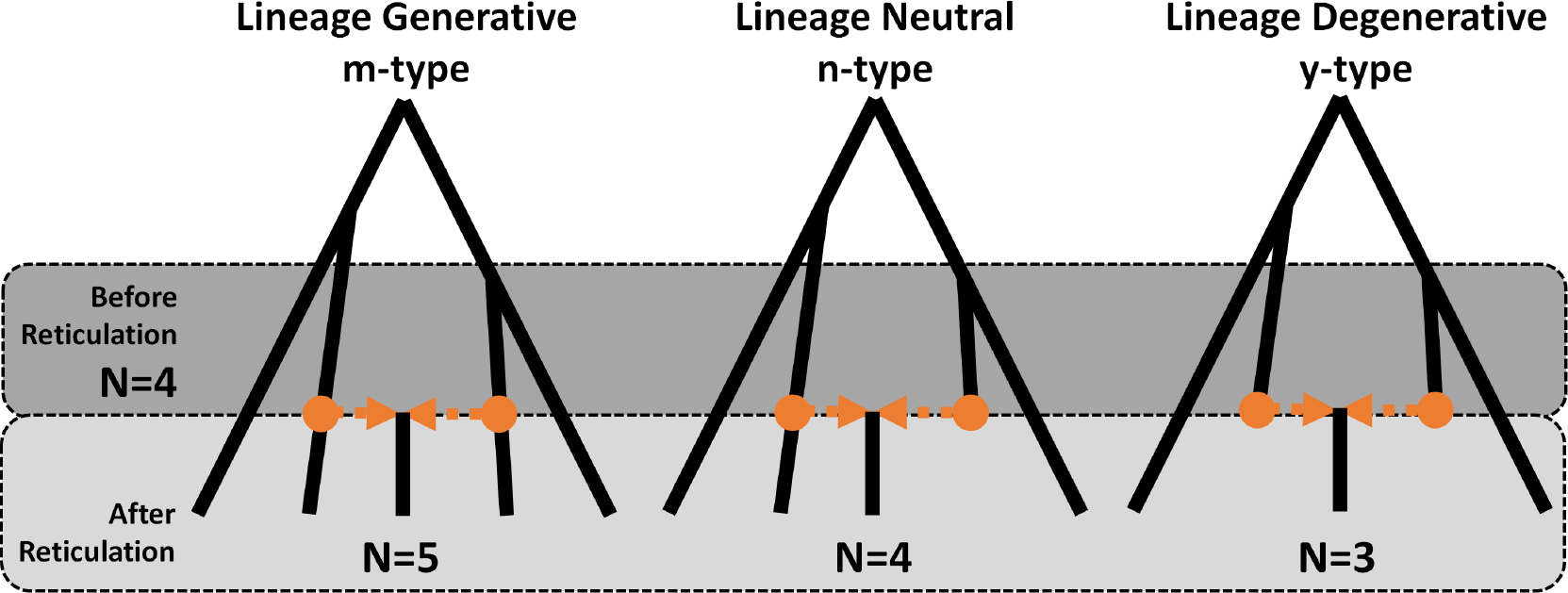
Types of Reticulation. Orange circles denote the parental nodes leading to the reticulate node. The dark grey box delineates the period before the reticulation event when the number of lineages is *N* = 4. The light grey box highlights the period after reticulation. Lineage generative hybridization (m-type) occurs when a reticulation event results in a gain of one lineage (*N* = 5). Lineage neutral hybridization (n-type) results in a net zero change in the number of lineages (*N* = 4), and degenerative hybridization (y-type) reduces the number of lineages by one (*N* = 3).

| Reticulate Pattern | Possible Biological Processes |
| --- | --- |
| **Lineage Generative** | • Hybrid speciation<br>• Allopolyploidy |
| **Lineage Neutral** | • Genetic swamping with one parental lineage<br>• Introgression<br>• Lateral gene transfer<br>• Lineage generative event with one unsampled parental lineage |
| **Lineage Degenerative** | • Genetic swamping with both parental lineages<br>• Lineage generative event with both parental lineages unsampled<br>• Lineage neutral event with one unsampled parental lineage |

The type of reticulation also affects the topological properties of phylogenetic networks (Janssen and Liu 2021). Phylogenetic networks can be grouped into a multitude of classes based on their topological features (see Box 2 and Zhang 2019; Kong et al. 2022). Understanding the topological properties of phylogenetic networks is exceptionally important because of the scalability challenges associated with inference (Hejase and Liu 2016; Elworth et al. 2019). Some network classes have useful identifiabilty results (Willson 2010b; Pardi and Scornavacca 2015; Solís-Lemus and Ané 2016; Francis and Moulton 2018; Erdős et al. 2019), unique encodings (Van Iersel and Moulton 2014; Linz and Semple 2020; Semple and Toft 2021), or polynomial-time algorithms for reconstruction (Erdős et al. 2019; Murakami et al. 2019; Semple and Toft 2021). Consequently, some methods will assume *a priori* that the underlying network belongs to a given class. For example, both the algorithms of Bordewich and Tokac (2016) and Wawerka et al. (2022) assume that networks are tree-child, while the software packages SNaQ (Solís-Lemus and Ané 2016) and NANUQ (Allman et al. 2019) assume that networks are level-1 (see Box 2 for definitions of network classes). Thus, to ensure methodological assumptions are met, it is important to understand the complex topological landscape of phylogenetic networks.

Some analytical results exist for network classes; however, they are currently only applicable to special cases of the birth-death-hybridization process and may be difficult to generalize (Bienvenu et al. 2022). Nevertheless, simulation has been an essential technique for characterizing the distribution of phylogenetic networks generated from various processes (Arenas et al. 2008; Janssen and Liu 2021). Through simulation, researchers found that increasing the number of hybridizations on a network of fixed size decreased the proportion of networks that belonged to certain classes (e.g., tree-based and tree-child; Janssen and Liu 2021). Additionally, for branching processes—including a special case of the birth-death-hybridization process—the type of reticulation also affected the topological properties of simulated networks. However, when comparing types of reticulation, a different simulation tool was used for each type, making it difficult to distinguish the effects of the reticulation type from artifacts of the generators themselves. Lastly, simulating to a fixed number of taxa and hybridizations obfuscates the role of hybridization in lineage diversification. The birthdeath-hybridization process offers a complimentary approach for analyzing specific properties of phylogenetic networks. By having all three reticulation types as a part of the model, we can directly assess their effects on topological properties and lineage growth of phylogenetic networks.

In this study, we examine and summarize the distribution of phylogenetic networks generated by the birth-death-hybridization process. Specifically, we used a unified modeling framework to consider different assumptions about the type reticulation events. Additionally, we modeled a genetic distance dependence on hybridization by decreasing the rate of hybridization linearly based on the relatedness between two species. To understand the impact of hybridization on lineage diversification in phylogenetic networks, we simulated the average of lineages through time. We also simulated phylogenetic networks and characterized the proportion of phylogenies that belong to common network classes. We simulated phylogenies with incomplete lineage sampling and genetic distance-dependent hybridizations to consider how important systematic and biological processes might affect the properties of generated phylogenetic networks.

Our work explores the space of phylogenetic networks generated under the birth-death-hybridization process and assesses how different assumptions about the process of gene flow influence this distribution. Developing an intuition for how hybridization creates and shapes diversity will be valuable for selecting and fitting appropriate macroevolutionary models. Lastly, understanding the distribution of topological properties from a biologically relevant process can give insight into how often network class assumptions are violated in empirical systems, potentially biasing inferences and steering methods development.

## 2 Methods

### 2.1 Phylogenetic Network Properties

Phylogenetic networks are similar to phylogenetic trees but have additional reticulate nodes and edges representing genetic material crossing species boundaries. Although these nodes can represent specific biological processes such as lateral-gene transfer or introgression, the network does not distinguish which process occurs. Though, for consistency with the formal descriptions of the birth-death-hybridization process (Morin and Moret 2006; Woodhams et al. 2016; Zhang et al. 2018), we, at times, refer to reticulation events as hybridization nodes. Like phylogenetic trees, networks can be rooted or unrooted. In this study, we consider rooted phylogenetic networks. Formally, a phylogenetic network is a directed acyclic graph that contains four node types:

- *root* : A node of in-degree 0 and out-degree 1 if the process begins at the root or out-degree 2 if the root represents the most recent common ancestor of the process.
- *tree*: A node of in-degree 1 and out-degree 2. These nodes typically represent speciation events.
- *hybrid* : A node of in-degree 2 and out-degree 1. Hybrid nodes denote reticulation events.
- *leaf* : Nodes of in-degree 1 and out-degree 0 that can represent either extant lineages that survived until the present or they can denote an extinction event.

A directed edge from nodes *u* to *w* is denoted as *(u,w)*. We further describe *(u,w)* as a hybrid edge if *w* is a hybrid node. Hybrid edges additionally have weights (*γ* and 1 − *γ*) that are used to describe the proportion of genetic material that contributes to the hybrid node. Hybrid nodes are considered stacked if the child of a hybrid node is another hybrid node (*i.e*., there exists an edge *(x,y)* where both *x* and *y* are hybrid nodes). We restrict our analyses to identifying tree-based, fold-unfold stable (FU-stable), tree-child, normal, and level-*k* networks (Box 2).

**Box 2: Description of Network Classes**

Phylogenetic network classifications describe the topological properties of a given phylogenetic network. In this study, we focus on five different classifications:

- **Level-***k*: Any biconnected component of the network having at most *k* hybrid nodes (Jansson and Sung 2006).
- **Tree-based**: A phylogenetic network that can be constructed by starting with a phylogenetic tree and sequentially adding reticulation edges (Francis and Steel 2015). Non-tree-based networks are identified by locating a zigzag pattern that connects reticulate nodes (Zhang 2016).
- **Fold-unfold stable (FU-stable)**: A network that can be ‘unfolded’ into multi-labeled trees (MUL-trees) and then ‘refolded’ (see Huber and Moulton 2006) back into itself is FU-stable. Fold-unfold stability occurs when reticulate nodes are not stacked and when no two tree nodes have the same set of child nodes (Huber et al. 2016).
- **Tree-child**: A tree-child network is a phylogeny where all internal nodes have at least one non-reticulate child node (Cardona et al. 2008).
- **Normal**: tree-child networks that contain no directed ‘shortcut’ edges that directly connects two nodes (*u, v*) given that there exists another directed path from node *u* to *v* (Willson 2008, 2010b).

The latter four classes have the following nested hierarchy:

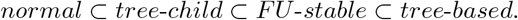

Any nested class will also have the topological properties of the all superceding classes (e.g., a tree-child network is also FU-stable and tree-based). Figure **??** gives positive and negative examples of the classifications, but also see Kong et al. (2022) and the webpage “ISIPhyNC (Information System on Inclusions of Phylogenetic Network Classes)”^*a*^ for more information on each classification.

^*a*^http://phylnet.univ-mlv.fr/isiphync

### 2.2 Birth-Death-Hybridization Process

We use a birth-death-hybridization process (Morin and Moret 2006; Woodhams et al. 2016; Zhang et al. 2018) as a generating model for phylogenetic networks. This process has exponentially distributed waiting times between speciation, extinction, and hybridization events. We consider rates of speciation (*λ*) and extinction (*μ*) to be rates on each lineage, while the rate of hybridization (*ν*) is on each lineage pair. That is, when the process has *N* lineages, the rates of speciation, extinction, and hybridization are *Nλ, Nμ*, and 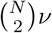, respectively. There are three types of hybridization events: lineage generative hybridization, lineage neutral, and lineage degenerative (Box 1). We use the thinning property of a Poisson process to break the hybridization rate (*ν*) into separate rates for each type of hybridization: *ν* = *ν*_+_ + *ν*_0_ + *ν*_−_, where the lineage generative hybridization rate is denoted as *ν*_+_, lineage neutral hybridization as *ν*_0_, and lineage degenerative hybridization as *ν*_−_. We refer to the possible state space—all linear combinations of *ν*_+_, *ν*_0_, and *ν*_−_ for a given hybridization rate (*ν*)—as the hybridization rate simplex, which we represent visually using a ternary plot (see example in Fig. 2).

**Figure 2:**
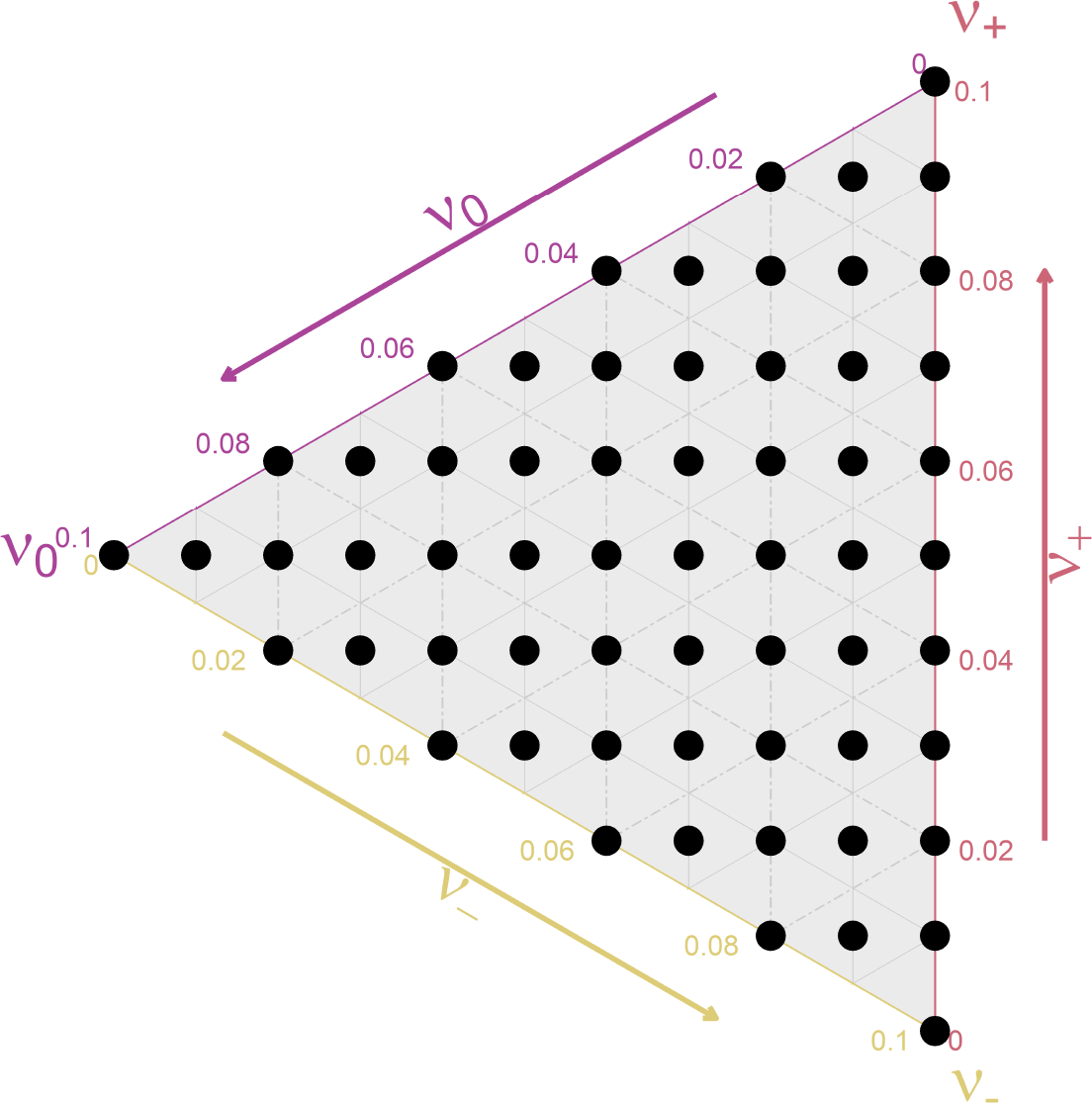
Ternary plot with the state space for the combination of hybridization-type rates for *ν* = 0.1. Moving towards a corner corresponds to an increase of the rate for one of the hybridization types: the top corner (*ν*_+_) denotes the lineage generative rate, the bottom corner (*ν*_−_) denotes the lineage degenerative rate, and the leftmost corner (*ν*_0_) denotes the lineage neutral rate. Each dot represents one parameter combination used when simulating across the hybrid rate simplex (Table S1). See Section 2.4.2 for details about the simulation procedure.

At a given hybridization event, we denote the genetic contribution of each parental lineage as *γ* and 1 − *γ*. Each hybridization event has its own associated *γ* that can be drawn from any distribution on [0, 1]. We draw the inheritance proportions *γ* from a *Beta*(10, 10) distribution, creating a symmetric distribution centered on equal inheritance proportion of *γ* = 0.5.

In some of our simulations, hybridization events are more likely to occur if the the two parent lineages are closely related. To accomplish this, we used the approach of Woodhams et al. (2016) for genetic distance– dependent hybridization. Under this model, hybridization events are proposed with rate *ν* and successfully occur on the phylogeny with a probability that is proportional to the genetic distance between the two hybridizing taxa (*d*_*ij*_):

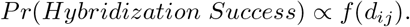

We assume a strict molecular clock where the genetic distance between two taxa is taken to be the summation of distinct branch lengths between each lineage and their most recent common ancestor. Moreover, in the case of hybrid taxa, the distance is a weighted summation across paths (see Woodhams et al. 2016).

There are many ways to model the relationship between genetic distance and the probability of hybridization (See Woodhams et al. 2016). For example, the Dobzhansky-Muller incompatibility model predicts that hybrid incompatibilities will accumulate at an accelerating rate (Orr and Turelli 2001). Here, we use a linear decay function to relate the probability of hybridization success to genetic distance:

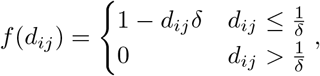

where *δ* represents the strength of the genetic distance dependence. A value of *δ* = 0 makes hybridization events independent of genetic distance, while higher values of *δ* make hybridization primarily occur between more closely related taxa.We can also see that, for genetic distances where *d*_*ij*_ is greater than the inverse of the dependence strength 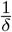, it is not possible for hybridization to occur.

Note that the generating process described above and our simulations differ slightly due to the stopping conditions we applied in our simulations. We focus on extant phylogenies during simulation and condition the process on survival of more than one taxon. Further, for practical purposes, we limit the run time of simulations due to the potential for some simulation replicates to never reach the specified ending age in finite time (see Section 2.4.2).

### 2.3 Diversification Dynamics

Measuring the rate of lineage accumulation over time allows us to compare the outcomes of different models. We examine how the diversification dynamics change both with the number of lineages and when we change our assumptions about hybridization to be either be primarily lineage generative or degenerative. Additionally, we show that, for certain parameter values of the birth-death-hybridization process, the process ‘explodes,’ reaching an infinite number of lineages in finite time.

#### 2.3.1 Diversification Rate

In a non-reticulate birth-death process, the net diversification rate is simply *λ* − *μ* (e.g., Nee 2006; Morlon et al. 2010). However, under the birth-death-hybridization process, net diversification has a different interpretation since hybridization events also can cause the number of lineages to increase or decrease. Further, because hybridization rates scale quadratically with the number of lineages, net diversification cannot be computed as the difference between all lineage adding and lineage deleting rates, *i.e*., (*λ* + *ν*_+_) − (*μ* + *ν*_−_). Instead we can compute the overall diversification rate for a process with *N* lineages as:

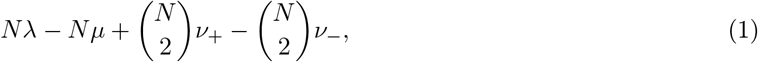

or equivalently:

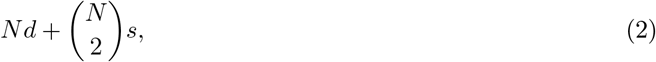

where *d* = *λ* − *μ* and *s* = *ν*_+_ − *ν*_−_. Thus, we interpret *d* as the per-lineage diversification rate and *s* as the lineage-pair diversification rate due to hybridization. The overall diversification rate is a function of the number of species but allows us to more directly compare the effects of speciation, extinction, and hybridization on lineage accumulation. From this perspective we can see that, as the number of lineages increases, hybridization quickly becomes the dominant force in diversification (Fig. 3a). The overall diversification rate grows rapidly when *s* is positive but actually becomes negative with a large number of taxa and when *s <* 0 (Fig. 3b). However, when considering a negative value of *s* and lineages through time, the overall diversification rate approaches zero as the negative effects from hybridization match the per-lineage diversification.

**Figure 3:**
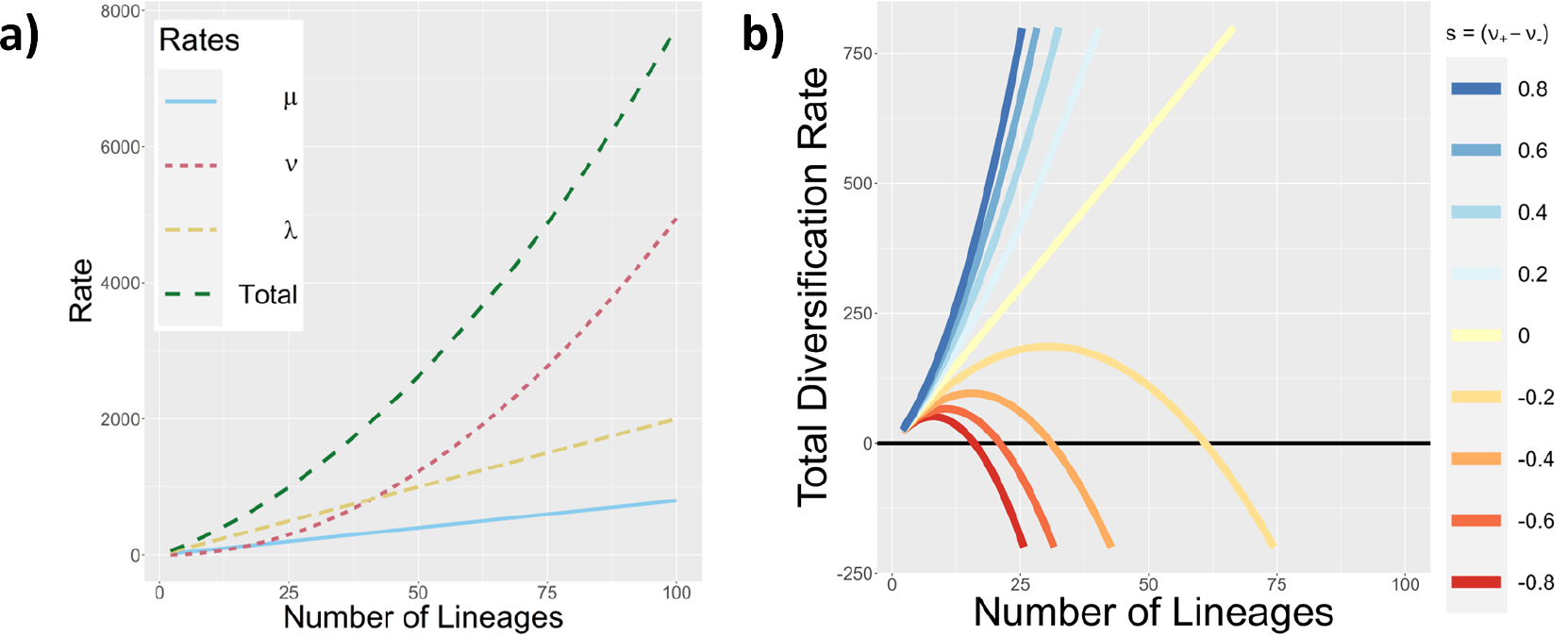
The effect of hybridization type on diversification dynamics. For all panels, we assume a speciation rate of *λ* = 20, extinction rate of *μ* = 8, and a hybridization rate of *ν* = 1. **a)** The rate of speciation(*λ*), extinction(*μ*), and hybridization(*ν*) events as the number of lineages increase. Panel **b)** shows the overall diversification rate 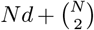s as a function of the number of lineages *N*.

#### 2.3.2 Explosion of the Birth-Death-Hybridization Process

Birth-death processes with sufficiently high birth rates can result in ‘explosive’ growth where an infinite number of events happen in finite time. We show that under certain conditions the birth-death-hybridization model exhibits ‘explosive’ growth. The rate of hybridization events scale with the total number of species pairs, creating a quadratic feedback term. Thus, when the rate of lineage generative hybridization exceeds the rate of lineage degenerative hybridization, new lineages tend to be added fast enough to induce explosion.

Specifically, we show that the process explodes when *λ >* 0, *μ* ≥ 0, and *s >* 0 (*i.e*., *ν*_+_ *> ν*_−_). If *ν*_+_ *>* 0 and *ν*_−_ = 0, the birth-death-hybridization process becomes a special case of the birth-death process with combinatoric innovation and is known to explode in finite time (Steel et al. 2020). In this section we will assume that *ν*_−_ *>* 0 and show that explosion occurs when *s >* 0. For now, we will also assume that *λ > μ*. In the supplementary materials we will relax this assumption and provide a proof for explosion. Theorem 3.2.2 from Anderson (1992) (equivalently theorem 2.2 of Bansaye and Méléard 2015) states that a birth-death process almost surely has an infinite lifetime if and only if the following sum diverges

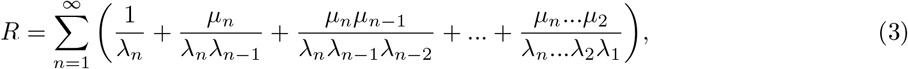

where *λ*_*n*_ and *μ*_*n*_ are the birth and death rates when the process has *n* lineages. Therefore, we can show the process almost surely has a finite lifetime (*i.e*., explodes) if and only if the series instead converges. For the birth-death-hybridization process, both speciation and lineage generative hybridization add new lineages while extinction and lineage degenerative hybridization remove lineages, giving birth rate:

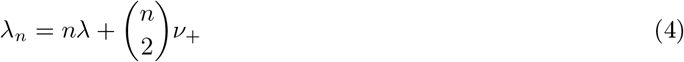

and death rate:

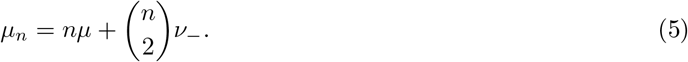

Defining 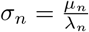 gives:

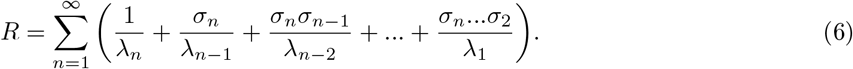

The sum can be written out as follows:

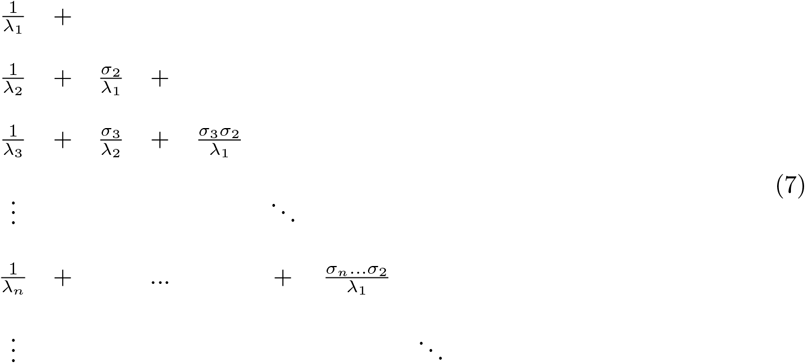

and *R* can be rewritten to be the sum along each diagonal *S*_*n*_, collecting each term with 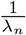 into its own summation:

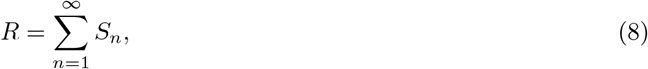

where:

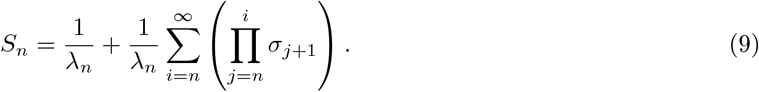

We can show that *S*_*n*_, and consequently *R*, converge by considering the sum: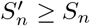

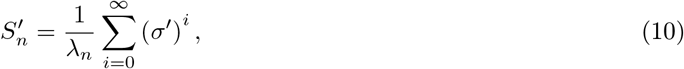

where *σ*^*′*^ is the supremum of *σ*_*n*_ when *n* ≥ 1:

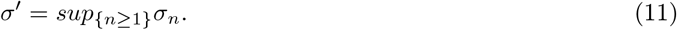

Since *λ* − *μ* and *s* are positive, *σ*_*n*_ *<* 1 for all *n*. Consequently *σ*^′^ *<* 1, making 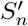 a geometric series that converges on 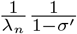. We can then define *R*^*′*^ to be the summation of all 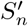:

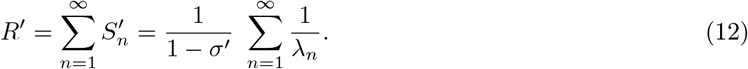

Here *R*^*′*^ *<* +∞ because *λ*_*n*_ grows quadratically with *n* when *ν*_+_ *>* 0. By direct comparison, we get *R < R*^′^. We can then conclude that *R <* +∞, meaning that the process can explode in finite time.

### 2.4 Birth-Death-Hybridization Process Simulations

In this study, we sought out to assess how different assumptions about hybridization affect the distribution of phylogenetic networks under a birth-death-hybridization process. We simulated both the number of lineages over time and phylogenetic networks to evaluate properties and expectations that are difficult to compute analytically under a birth-death-hybridization process. All simulations were conducted on a Dell XPS 15 with an i7 processor.

#### 2.4.1 Simulating Lineage Diversification over Time

To understand the accumulation of lineages under the birth-death-hybridization process, we simulated the process over time using the R package GillespieSSA (avaliable on CRAN; Pineda-Krch 2008). This package allows for efficient simulation of the number of lineages in two ways: (1) it can simulate under the birthdeath-hybridization process without having to simulate and record the structure of the phylogeny and (2) each simulation records the number of lineages at multiple time points, not just at the end of the simulation. All simulations began with two lineages at *t* = 0 and were allowed to proceed until reaching the stopping time *t* = 10. We used constant rates of speciation (*λ* = 4), extinction (*μ* = 3), hybridization (*ν* = 0.2), and assumed no genetic distance dependence (*δ* = 0). Since lineage neutral hybridization does not change the number of lineages in the process, we assumed a lineage neutral hybridization rate of zero (*ν*_0_ = 0). To change the lineage-pair diversification rate *s*, we modulated the values of lineage generative and degenerative hybridization between simulations to assess how different lineage-pair diversification rates *s* change the diversification dynamics. Since positive values of *s* can lead to an infinite number of lineages (see Section 2.3.2), we limited each simulation to run for 4 seconds. For each value of *s*, we simulated until we collected 5,000 replicates that survived until *t* = 10 or 5000 replicates that reached the timeout time of 4 seconds.

#### 2.4.2 Simulating the Distribution of Phylogenetic Network Classes

Phylogenies were simulated using the R package SiPhyNetwork (avaliable on CRAN; Justison et al. 2023). Simulations were conditioned on survival to the present with at least two extant lineages prior to sampling and to have completed in less than 10 seconds. The latter condition is to prevent simulation replicates from spending an infinite amount of time approaching the stopping time without ever reaching it. We expect this behavior for positive lineage-pair diversification rates (*s >* 0).

All simulations began with two lineages at *t* = 0, representing the most recent common ancestor at the root, and were allowed to proceed until reaching the stopping time at *t* = 1. We used constant rates of speciation (*λ* = 4), extinction (*μ* = 2), and hybridization (*ν* = 0.1). For each specific scenario and parameter combination (Tables S2, S3, and S4) we generated 20,000 replicate phylogenetic networks.

#### 2.4.3 Simulating Across the Hybridization-Rate Simplex

To assess how hybridization type affected the properties of simulated phylogenies, we used 61 different linear combinations of rates for the hybrid types *ν*_+_, *ν*_0_, and *ν*_−_. The specific parameter combinations are visualized in Fig. 2 and listed in Table S1.

For each combination of hybridization-type rates, we used two different sampling strategies of extant-only sampling and complete sampling. First, we considered complete phylogenies with all extant and extinct taxa included in the phylogeny. For these complete phylogenies we had two scenarios where hybridization was either genetic distance–dependent (*δ* = 0.5) or independent of genetic distance (*δ* = 0). Next, we sampled extant-only phylogenies by pruning extinct taxa from the phylogeny. For the extant-only phylogenies, we considered both complete extant sampling (*sampling fraction* = 1) and incomplete sampling where a constant fraction of extant taxa were sampled (*sampling fraction* = 0.75).

We additionally simulated phylogenies to isolate the effects of genetic-distance dependence and incomplete sampling on the class membership of networks. We used a constant rate for each hybridization type 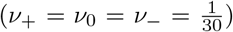 while separately varying the strength of genetic-distance dependence and amount of incomplete lineage sampling. For incomplete lineage sampling, we simulated extant only phylogenies and used sampling fractions that ranged from *ρ* = 0.25 to *ρ* = 1.0. For genetic distance dependence, we simulated complete phylogenies with both extinct and extant taxa while varying the strength of distance dependence from *δ* = 0 to *δ* = 2.

### 2.5 Analysis of Simulated Data

For each replicate of the lineage diversification simulations, we recorded the number of lineages through time. We then reported the average number of lineages over time for each lineage-pair diversification setting for *s* when conditioning on survival at each time point. We analyzed the simulated phylogenetic networks using a combination of summary statistics and network classifications to characterize the distribution of phylogenetic networks under the birth-death-hybridization process. For the replicates of each simulation scenario, we recorded the total number of taxa, total number of hybridization events, reticulation density 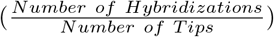, observed proportion of each hybridization type, and the level of the network. Lastly, we assessed whether phylogenies belonged to the tree-based, FU-stable, tree-child, or normal classes (Box 2).

## 3 Results

Our aim—through both analytical computation and simulation—was to understand and characterize the properties of phylogenetic networks generated from the birth-death-hybridization process. We also assessed how our assumptions about gene flow and sampling affected this distribution. We present and outline our key findings below.

### 3.1 Lineage Diversification over Time

#### The shape of lineage accumulation took different forms depending on the relative rates of lineage-generative and lineage-degenerative hybridization (Fig. 4)

When *ν*_+_ *> ν*_−_, the lineagepair diversification was positive, and the average number of lineages exhibited hyper-exponential growth. In fact, all simulation settings with *s >* 0 reached 5,000 timeouts before reaching 5,000 surviving simulations.

This result was consistent with the explosive behavior that can occur when *s >* 0, where too many events— possibly infinitely many—would occur before the simulation could finish. When conditionined on survival, *ν*_+_ *< ν*_−_ resulted in a negative value for *s*, the average number of lineages approached a constant number (Fig. 4). The effects of positive diversification via speciation and extinction (*i.e*., *Nd*) were neutralized by negative diversification due to hybridization (*i.e*., 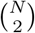) and resulted in a net-zero total diversification (see Eq. 2). Interestingly, when not conditioned on survival and *s <* 0, the average number of lineages decreased over time (Fig. S2). Any time the process reached a large enough population, lineage degenerative hybridization acted a diversity dependent mechanism that inhibited growth. Then over a long enough period of time, the process would eventually run to extinction due to the stochastic nature of events, causing the decline in the average number of lineages.

**Figure 4:**
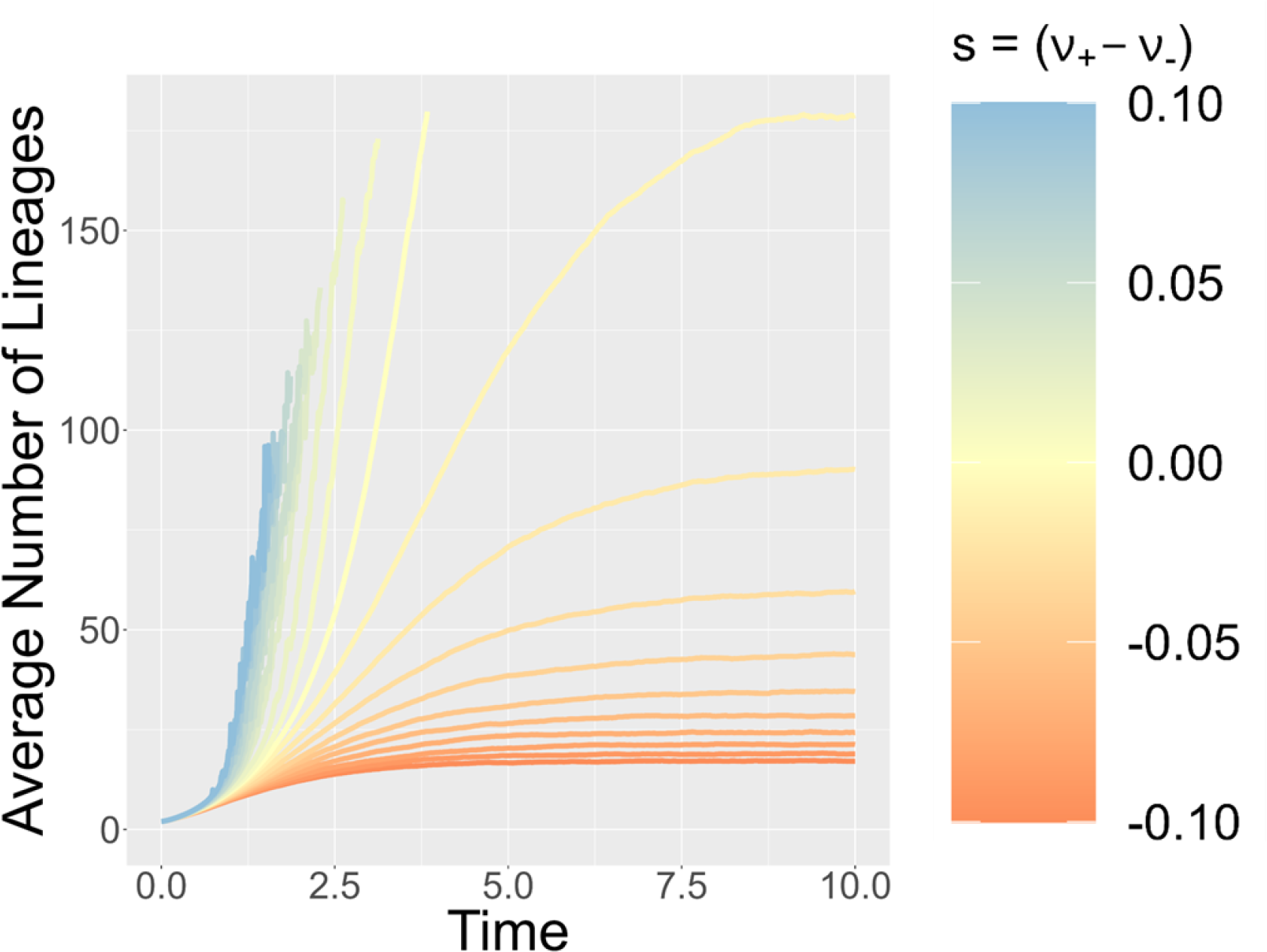
The average number of lineages through time with respect to *s* when conditioned on survival. We assume a speciation rate of *λ* = 4, extinction rate of *μ* = 3, a hybridization rate of *ν* = 0.2, and a lineage neutral hybridization rate of *ν*_0_ = 0. Higher lineage-pair diversification (*s*) corresponds to a darker blue while low values of *s* are denoted with a dark orange. Lines were only drawn for time points that had at least 4,500 replicates that had survived to that time point and had not timed out.

### 3.2 Phylogenetic Network Simulation

We generated 20,000 phylogenies that met the criteria defined in Section 2.4.2 for each parameter combination (Tables S3 and S4). Across all completely sampled simulations without genetic-distance dependence, we discarded 339,054 simulation replicates (21.7% of total attempts) that went extinct or yielded only one extant taxon. We discarded 2,233 replicates (0.14% of total attempts) due to the simulation exceeding the 10 second stopping condition. A higher proportion of simulations timed out as *s* increased. Increasing the stopping time to 20 seconds for the settings with the highest lineage-pair diversification (*s*) did not have a large affect on the number timeouts (1,074 timeouts; 0.13% of simulations), indicating that we were likely able to capture the distribution of networks with little bias. Simulations rejected for timing out may have never finished or were outliers in terms of numbers of extant lineages.

### 3.3 Simulating Across the Hybrid Rate Simplex

#### Hybridization type affected the diversification dynamics of simulated phylogenies (Fig. 5)

In accordance with analytical expectations, simulations with higher lineage-pair diversification rates (*s*) had an increased number of taxa on average, a result that is consistent across all sampling scenarios (complete, complete with genetic distance–dependent hybridization, extant-only networks, and extant-only networks with incomplete sampling). We also observed the same increasing pattern with respect to *s* for the number of hybridization events and reticulation density. Interestingly, however, extant-only networks had higher reticulation densities than complete networks despite having fewer reticulations (Fig. 5; Table S5). Though all extinct lineages are not observed for extant-only sampling, reticulations are only lost on the phylogeny when all the hybrid descendants fail to get sampled. This becomes increasingly difficult for highly intertwined networks with many reticulations. Further, due to incomplete sampling among some parental lineages, extant-only networks had a higher proportion of lineage-degenerative and lineage-neutral hybridization than completely sampled networks with the same hybridization rates (Fig. S3).

**Figure 5:**
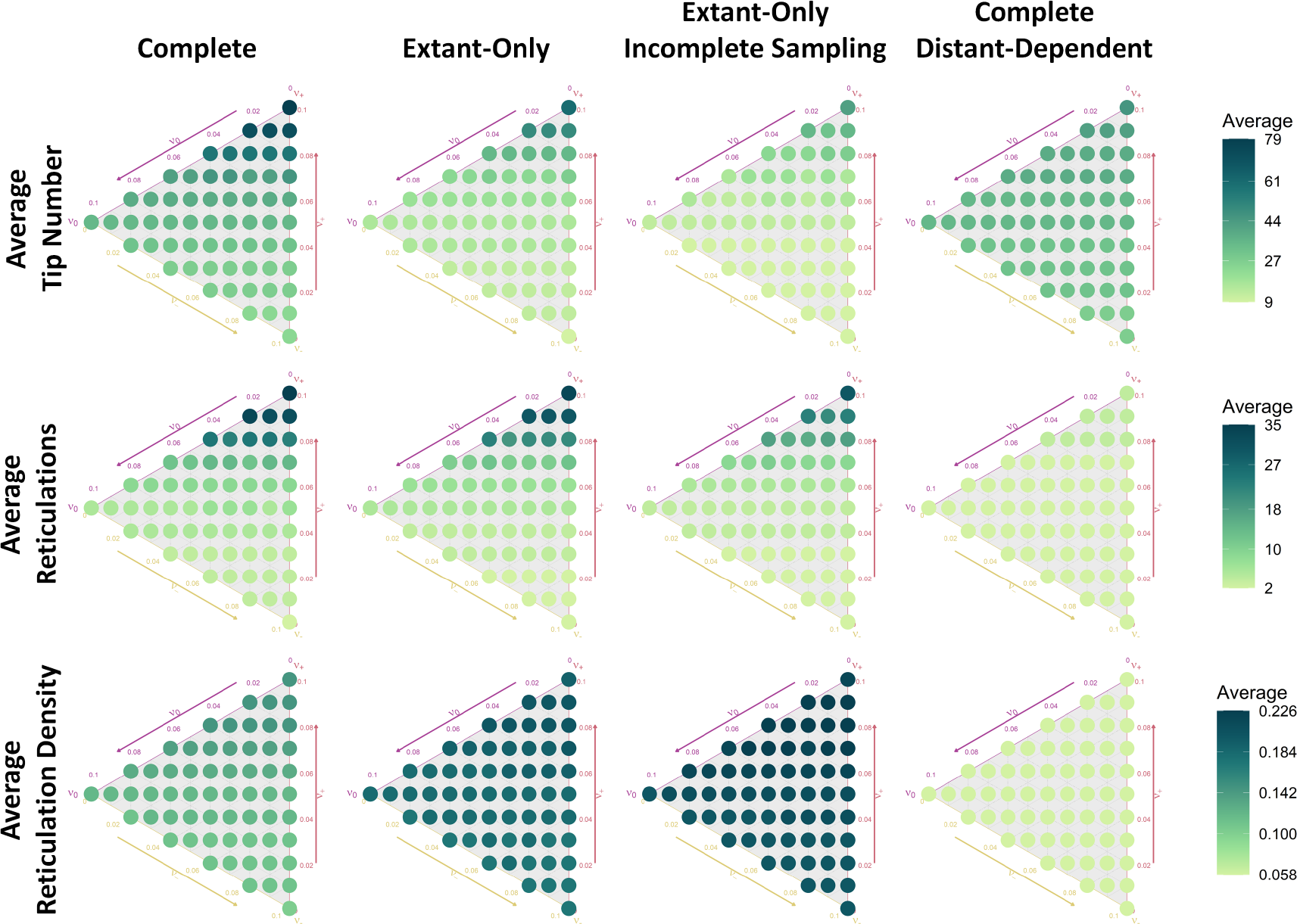
Diversification dynamics of phylogenetic networks simulated across the hybrid rate simplex. Each column shows one of the four simulation conditions used to simulate phylogenetic networks across the hybrid rate simplex (Section 2.4.2). The position of each point on the simplex indicates the lineage generative, lineage neutral, and lineage degenerative hybridization rates. Each row summarizes the number of tips, reticulations, and reticulation density 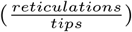 for each dataset. The color of each point is used to depict the average value of 20,000 replicates.

#### The proportion of networks belonging to each class was differentially affected across the hybrid rate simplex (Fig. 6; Table S6)

The more deeply contained classes within the nested hierarchy of tree-based, FU-stable, tree-child, and normal classes (see Box 2) have increasingly stringent topological requirements and, consequently, lower class proportions (e.g., a smaller proportion of networks were tree-child than tree-based). More deeply nested classes also showed a greater range of proportions across the hybrid simplex (Table S6). Notably, for completely sampled, extant-only, and incompletely sampled datasets, the region with high lineage-generative rates (*i.e*., high *s*) led to the lowest proportion of tree-based and FU-stable networks but resulted in relatively high proportions for tree-child and normal classes, thus indicating tree-based and FU-stable classes could be more sensitive to a high number of hybridization events than other classes. However, for genetic distance–dependent hybridization, the proportion of tree-based and FU-stable networks tended to decrease as the value of *s* decreased. Furthermore, for all simulation scenarios, the proportion of tree-child and normal networks decreased in regions with fewer hybridization events that consequently also had an increased proportion of lineage degenerative hybridizations (*i.e*., lower *s*), suggesting hybridization type is and important factor for these classes. When comparing between sampling conditions, class proportions were lower for the incompletely sampled datasets than those that were completely sampled (despite having fewer hybridization events). The complete dataset simulated under genetic distance–dependent hybridization had relatively few reticulation events, resulting in overall higher class proportions than other sampling conditions.

**Figure 6:**
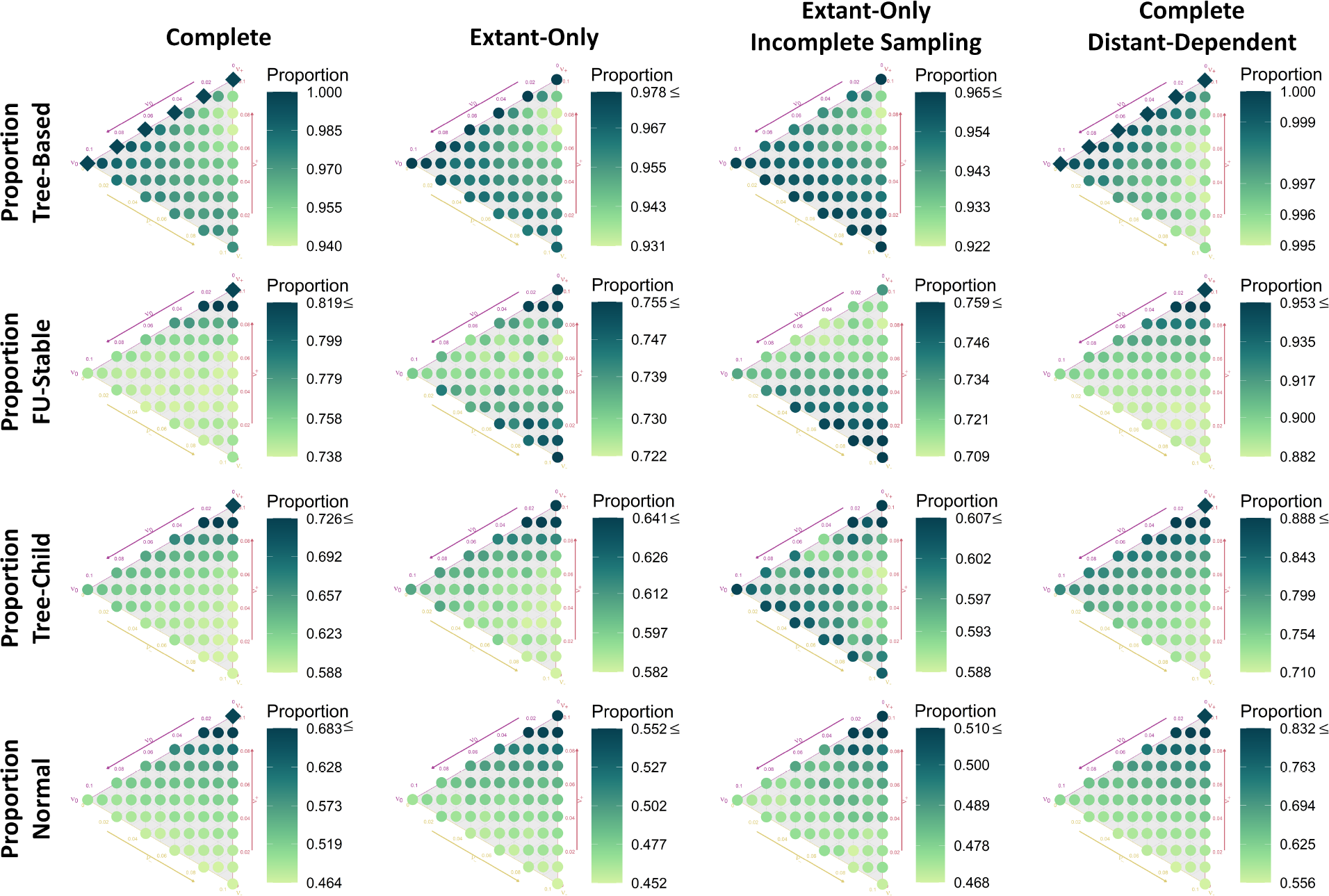
Class membership of simulated phylogenetic networks across the hybrid rate type simplex. Each column shows the simulation conditions used to simulate phylogenetic networks across the hybrid rate simplex (Section 2.4.2). The position of each point on the simplex indicates the lineage generative, lineage neutral, and lineage degenerative hybridization rates. Each row summarizes the proportion of phylogenies from a given class. The color of each point is used to depict the proportion of 20,000 replicates that belonged to the class, with a darker color representing a higher proportion. Colors are scaled for each class and dataset. Scales range from the minimum observed proportion to the 95^th^ percentile, and all values above the 95^th^ percentile are binned into the same color. Cases where all simulated networks belong to a class (proportion of 1) are indicated with a diamond.

Simulation replicates with only lineage-generative and neutral hybridization always produced networks with certain topological properties (dark diamonds in Fig. 6; Table S6). For completely sampled networks, all simulations with a lineage-degenerative rate of zero (*ν*_−_ = 0; axis with *ν*_0_) were tree-child. Similarly, at the corner of the hybrid rate simplex with only lineage generative hybridization (*ν*_+_ = *ν* = 0.1), all networks were tree-based, FU-stable, tree-child, and normal. Interestingly, the same parameterizations with extant-only or incomplete sampling did not lead to unity with respect to class membership. Unsampled parental lineages caused lineage generative and lineage neutral hybridizations to be observed as another type (e.g., Figures S1 and S3).

#### A large portion of reticulate phylogenies were not level-1 (Fig. S4)

Increasing the lineage-pair diversification rate (*s*) led to an increase in the average level of simulated networks and a higher proportion of level-2 (or higher) networks. Trends in the network level were strongly tied to the number of reticulations (Fig. 7). In most cases, the level of the network was the same as number of reticulations. However, there was slightly more variation in network level for those with few reticulations (Fig. 7b).

**Figure 7:**
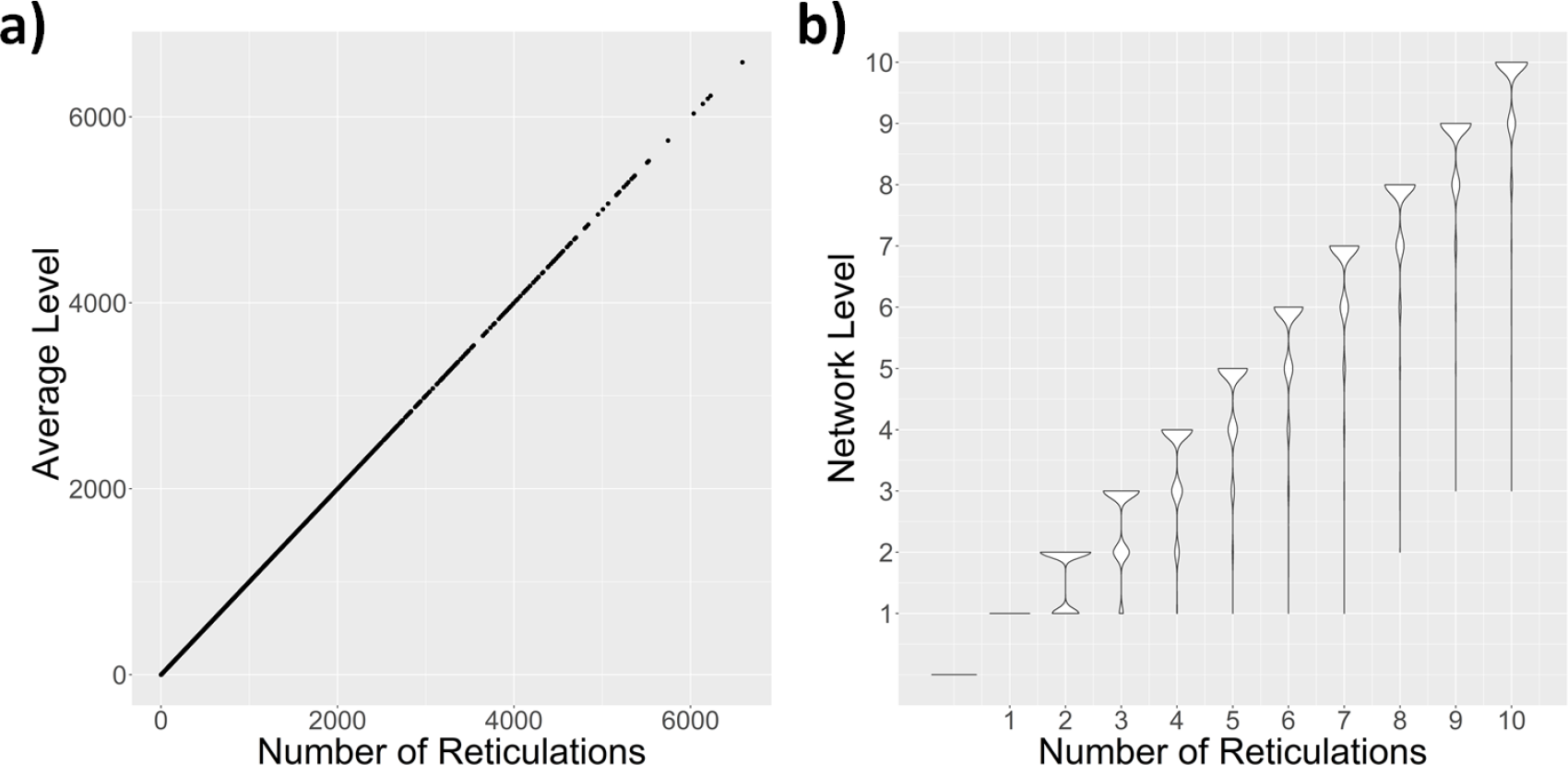
Completely sampled phylogenies summarized across the hybrid rate simplex by the number of reticulations. **a)** A scatterplot depicting a near one to one relationship between the number of reticulations and the average level of simulated phylogenetic networks. **b)** A violin plot showing the distribution of levels for 0 to 10 reticulations. Widths of the violin correspond to the proportion of networks with that level.

#### Increasing the effects of genetic-distance dependence and incomplete sampling decreased the number of reticulations on networks and consequently increased the class proportions of networks (Fig. 8)

Increasing the effect of incomplete sampling (*i.e*., decreasing the sampling fraction) only modestly affected the class proportions. However, increasing the effect of distance dependence (*i.e*., strength of distance dependence *δ*) more drastically increased class proportions, making nearly all networks tree-based and FU-stable. Higher class proportions can be explained by large decreases in the overall number of hybridization events and an increase in the proportion of phylogenies with no reticulation events, particularly for distance-dependent hybridization (Figures S6 and S7). For the incomplete sampling scenario, higher reticulation densities (Fig. S6) and changes in the observed hybridization types (Fig. S5) likely offset some of the effects of having fewer reticulations, explaining a more limited increase in class proportions. Interestingly, when looking on a per-reticulation basis, the opposite trend is observed for incomplete sampling. Decreasing sampling actually decreased class proportions for a given number of reticulations (Fig. 9). When looking on a per-reticulation basis for genetic distance–dependent hybridization, increased dependence strength (*δ*) is associated with higher proportions of tree-based and FU-stable networks but lower proportions of tree-child and normal networks.

**Figure 8:**
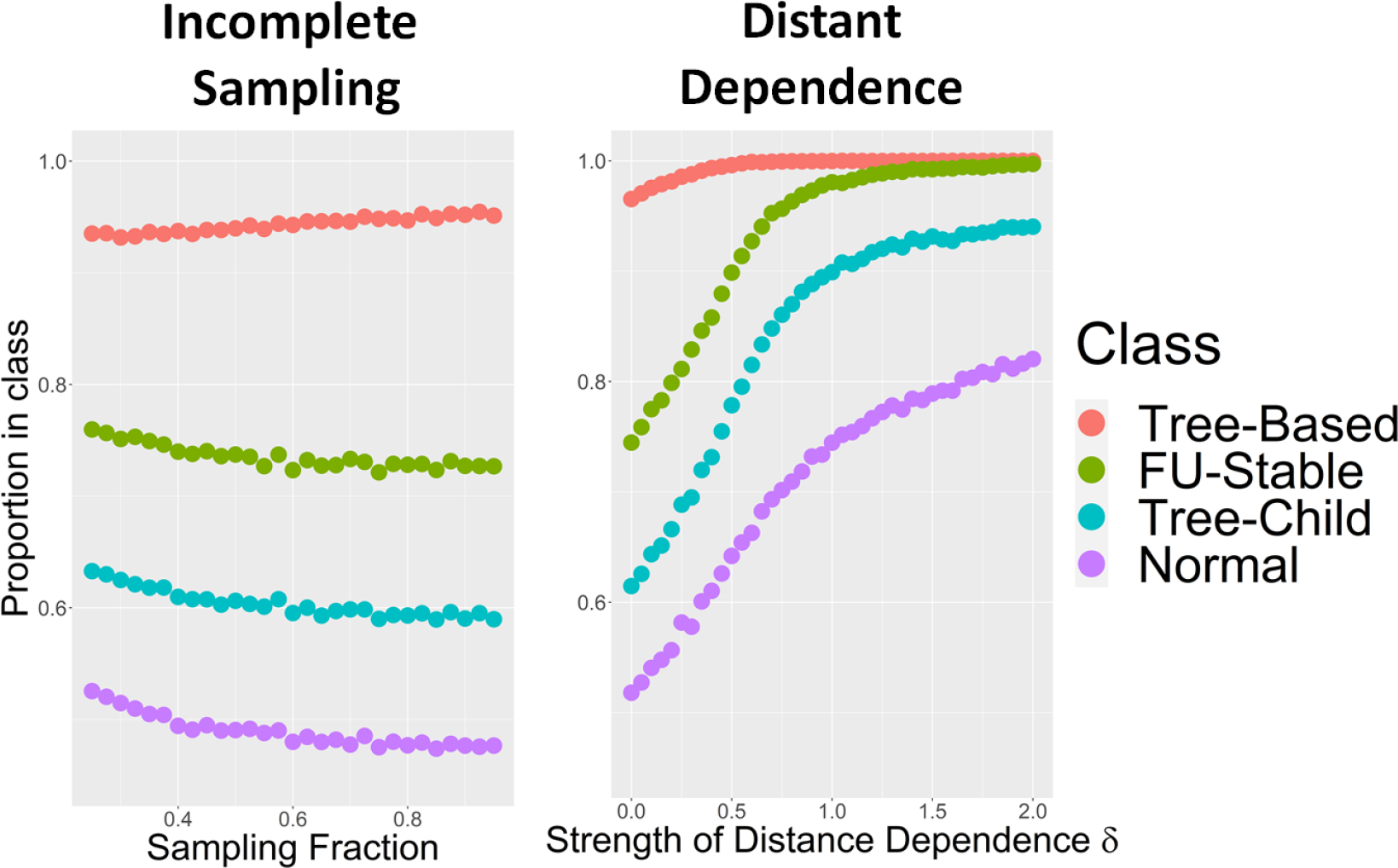
Proportion of simulated phylogenies that belong to network classes as a function of incomplete sampling and genetic distance dependence. Each point represents the class proportions observed from 20,000 simulated phylogenetic networks. The sampling fraction used in the incomplete sampling dataset represents the number of extant tips that are randomly sampled from the network. The value of *δ* used in the genetic distance-dependant dataset corresponds to the strength of dependence, with higher values of *δ* indicating that reticulation events primarily only occur between closely related lineages. Extant-only networks were used for the incomplete sampling dataset, while complete phylogenetic networks with extant and extinct species were used for the genetic distance-dependence dataset.

**Figure 9:**
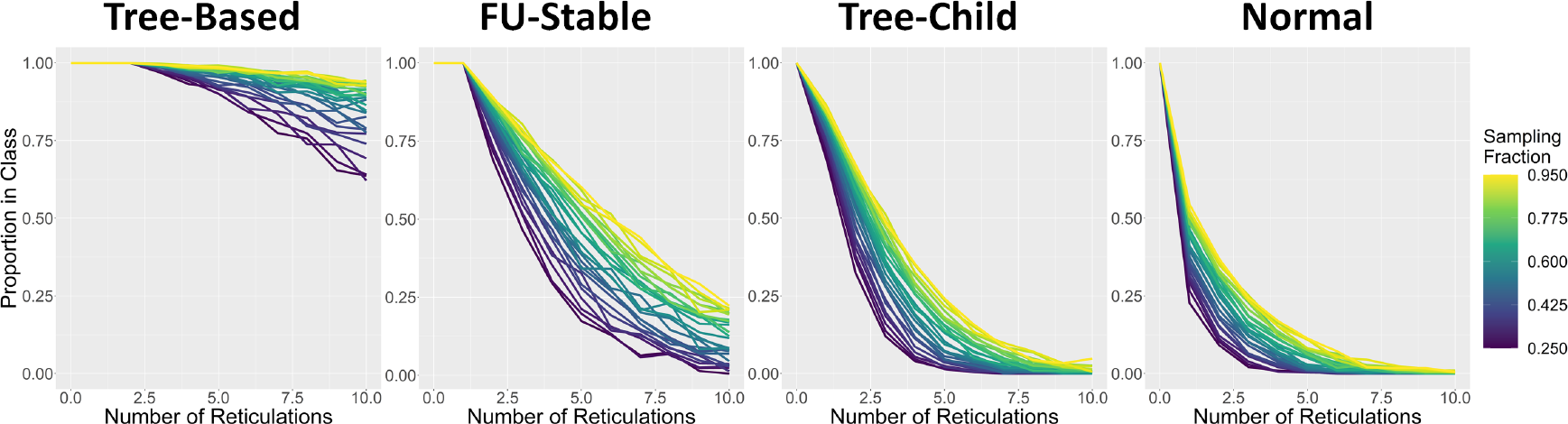
Class membership proportions under incomplete sampling as a function of reticulation number. Replicates are grouped and summarized by the number of reticulations and sampling fraction. The sampling fraction is the proportion of extant lineages sampled from an extant-only phylogenetic network. For each sampling fraction, 20,000 phylogenetic networks were simulated. The proportion in a class represents the proportion of networks with a given number of reticulation events to belong to that class. Points were only drawn when a given distance-dependence strength and number of reticulations had at least 30 replicates.

**Figure 10:**
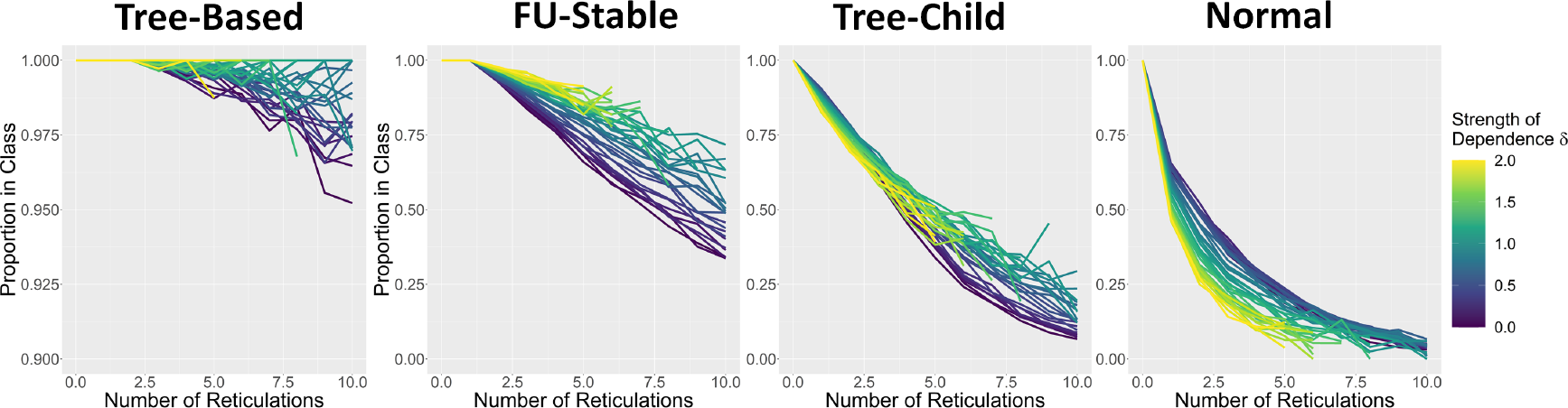
Class membership proportions under genetic-distance dependence and as a function of reticulation number. Phylogenetic networks are grouped by the simulated number of reticulations and strength of distance dependence. For each value of distance dependence (*δ*), 20,000 phylogenetic networks were simulated. The proportion in a class represents the proportion of networks with a given number of reticulation events to belong to that class.

## 4 Discussion

### 4.1 Diversification under Birth-Death-Hybridization

The birth-death-hybridization process is related to the logistic branching process (Lambert 2005) in that both allow for density-dependent feedback terms that affect growth; however, the birth-death-hybridization process allows for both positive and negative feedback based on the dominant type of hybridization. With the rate scaling by each lineage pair, hybridization can eclipse rates of speciation and extinction, resulting in highly reticulate histories. However, in nature, some biological systems exhibit wide-spread gene flow (Salzburger 2018; Edelman et al. 2019; Kozak et al. 2021; Esquerré et al. 2022; Suvorov et al. 2022), while it is much more limited in others (Solís-Lemus and Ané 2016; Morales-Briones et al. 2018; Zhang et al. 2018; Karimi et al. 2020). The lack of extensive reticulation in some empirical systems may be an artifact of the computational burden associated with estimating complex phylogenetic networks, particularly when applied to genome-scale data (Hejase and Liu 2016; Elworth et al. 2019). Empirical datasets may also result in a small number of inferred reticulation events due to the build-up of incompatibilities as lineages genetically diverge over time (Orr 1995; Mallet 2005; Gourbière and Mallet 2010). In fact, we showed that a linearly decreasing reticulation dependence on genetic distance had great propensity to reduce the number of events on simulated phylogenetic networks. However, there are a number of isolating mechanisms (see Mallet 2005; Soltis and Soltis 2009) that could create heterogeneity in the amount gene flow from system to system. Future work explicitly incorporating these mechanisms and modeling other genetic-distance dependencies into the birth-death-hybridization processes will be important in assessing their role in the diversification of admixed systems.

Solely changing assumptions about hybridization leads to phylogenies with considerable variation in size and pattern of diversification. The slowdown of lineage accumulation has been observed in numerous clades across the tree of life (Morlon et al. 2010). Several mechanisms have been proposed to explain slowing diversification (Moen and Morlon 2014), including density-dependent speciation rates (Rabosky and Lovette 2008) and protracted speciation (Etienne and Rosindell 2012). Under a birth-death-hybridization model where *s <* 0, hybridization acts as a diversity-dependent mechanism that reduces the overall degree of diversification and generates a pattern of slowed growth over time. In fact, estimating the likelihood of the phylogenetic network from a branching process is currently limited to a special case of the birthdeath-hybridization process where only lineage-degenerative hybridization occurs (*s <* 0; Zhang et al. 2018). Given that gene flow represents a broad set of processes with heterogeneous effects on diversification, it will be important to be able to estimate rates for each type of hybridization from the birth-death-hybridization process. Although available methods do not currently include extinction or estimate lineage neutral or lineage generative rates, for systems with lineage-generative reticulation, the hyper-exponential growth when *s >* 0 might prove helpful for estimating diversification dynamics. If these systems have ages calibrated by fossil or other geological data, it would be sensible to limit the parameter search space to regions of the hybrid-rate simplex that are not expected to explode to infinitely many species over the age of the clade or to invoke other biological processes that might limit such explosive growth.

### 4.2 Topological properties of Phylogenetic Networks

Two key results arise from previous simulation studies of topological profiles (Arenas et al. 2008; Janssen and Liu 2021): (1) as the number of reticulations increases, the proportion of generated networks that belong to certain classes (e.g., tree-child, tree-sibling) decreases and (2) when controlling for the number of reticulations, the type of reticulation affects the proportion of networks belonging to the same given classes. Our results are largely concordant with these conclusions. Though, when considering the birth-death-hybridization process, it creates a complex interaction of these factors; the reticulate events that are the best for some phylogenetic network classes (lineage generative; Janssen and Liu 2021) also lead to more lineages and consequently, more reticulations. We found that phylogenetic-network classes responded differently to these factors; tree-based and FU-stable network proportions were lower in the region of the hybrid rate simplex with a high number reticulations, primarily being lineage generative. Tree-child and normal classes had low proportions in the region that had few reticulation events but were mostly lineage degenerative. Thus, it is important to not only consider how each hybridization type affects the diversification and topological properties of systems but also to characterize how each network class individually responds to these factors.

Many network classifications require reticulations to be distant from one another. For example, networks will not be tree-child if reticulations are stacked, or networks are not tree-based if they have a specific zigzagging pattern that connects reticulations (Zhang 2016)—both only occur if reticulations have few nodes separating them on the phylogenetic network. Reticulation proximity makes it important to consider not only the number of hybridizations when considering topological properties but also their density with respect to the size a phylogenetic network. Both poor taxon sampling and not accounting for extinct lineages with fossil evidence has been shown to reduce the accuracy of phylogenetic inference (Heath et al. 2008a; Nabhan and Sarkar 2012; Warnock et al. 2020). Incomplete species sampling may also pose a challenge for estimating phylogenetic networks as the higher reticulation density of poorly sampled networks compared to those that are completely sampled may induce fewer topological properties that can inform phylogenetic estimates.

Both our simulations and those of Janssen and Liu (2021) with genetic distance dependence also highlight the importance of reticulation proximity. Gene flow reduces the observed divergence between species (Slatkin 1985; Leaché et al. 2014). When considering a fixed number of reticulation events, this creates a positive feedback loop for gene-flow events to occur locally on reticulate lineages, thus decreasing the proportion of networks with certain topological properties. However, these same distance-dependent mechanisms also reduce the overall number of gene-flow events, potentially mitigating the effects of increased locality. Further, we only considered a scenario where the hybridization rate decreases linearly with genetic distance, which has two potentially important characteristics: (1) the rate of hybridization may decrease slower than other biologically relevant mechanisms for modeling genetic-distance dependence (e.g., exponential or snowballing decay), and (2) after a certain threshold, the probability of two distant lineages hybridizing becomes zero. The former attribute may lead to excessive gene flow, which, due to the homogenizing effect of gene flow, would create a positive feedback loop that keeps the genetic distances between lineages low, resulting in more gene flow. The latter attribute effectively makes distant clades independent with respect to hybridization; this may be an undesirable effect if one is trying to model gene flow between distantly related lineages. Ultimately, if systems with distance-dependent gene flow (e.g., Chapman and Burke 2007; Tea et al. 2020; Barley et al. 2022) have an abundance of events, it may pose challenges to methods that make certain topological assumptions of the underlying phylogenetic network.

The proximity of reticulations to one another also explains why only lineage-generative or neutral hybridization (*i.e*., *ν*_−_ = 0) always produced networks of certain types and why incomplete or extant-only sampling eliminates this pattern (see lemmas 1-4 in Janssen and Liu 2021). Effectively, lineage-generative and neutral hybridization events create extra tree nodes during the reticulation event. These nodes can put space between reticulation nodes, making it more likely that a network meets certain topological criteria for class assignment.

While not all reticulate histories are tree-based and have an underlying tree as a backbone (Francis and Steel 2015), those generated from the our parameterizations of birth-death-hybridization process appear largely tree-based. The tree-based class is conceptually satisfying for phylogenetic networks by allowing researchers to think of reticulate evolution as phylogenetic trees with additional arcs. In fact, tree inference on reticulate systems has several uses and advantages over network inference (see Blair and Ané 2020). Though, inferring the treelike backbone of a network—if such a tree exists—should be done with caution because topology and divergence estimates can be biased (Leaché et al. 2014). Interestingly however, several recent methods attempt to quickly estimate phylogenetic networks by starting with phylogenetic trees and augmenting them into networks (Cao et al. 2019; Molloy et al. 2021).

Some relationships on phylogenetic trees are even impossible to infer (Slowinski 2001; Lewis et al. 2005), particularly for systems undergoing rapid speciation (Stanley et al. 2011; Suh 2016). Instead, polytomies can be included in an analysis to reflect the lack of phylogenetic resolution (Lewis et al. 2005; Kemp 2009). A similar discourse is occurring in the phylogenetic-network community with many studies finding only specific topologies identifiable and possible to estimate from gene tree topologies (Huber et al. 2015; Pardi and Scornavacca 2015; Solís-Lemus and Ané 2016; Francis and Moulton 2018; Gross and Long 2018; Erdős et al. 2019). Even when sequence alignments are used to alleviate identifiability issues, scalability and computational complexity remains a challenge for phylogenetic network inference. Pardi and Scornavacca (2015) proposed an interesting solution to the identifiability—and potentially scalability—problem: infer distinguishable canonical phylogenetic networks by effectively collapsing some reticulate events into polytomies. There are techniques capable of reconstructing the phylogenetic network if it is of a certain class and returns a reduced form network if not belonging to that class. Normal networks can be reconstructed in polynomial time from their set of displayed trees (Willson 2010a), and if the underlying network is not normal then the algorithm will return a reduced form normal network. Not all reticulations may be present in the reduced normal network, but these constructions have desirable mathematical properties and summarize the history into a simpler structure (Francis et al. 2021). Additionally, the folding operation of Huber and Moulton (2006) can reconstruct FU-stable networks from MUL-trees (Huber et al. 2016). If the underlying network is FU-stable then the method returns the true network, otherwise it produces a minimally reticulate network that unfolds into the same MUL-tree. Although estimating the MUL-tree is not a trivial task (Huber et al. 2008), MUL-trees and their implied FU-stable networks have been used to understand polyploid histories (Brysting et al. 2007). These types of approaches represent a shift from the fully resolved phylogenies that biologists have come to expect, but they have several appealing qualities: (1) they only return what is distinguishable, (2) if the true network has the right topological properties then it can still be recovered, and (3) canonical forms have a reduced topology search space with computationally efficient construction algorithms. Further, these approaches will still be applicable to the appreciable proportion of networks under the birth-death-hybridization process that do not have ideal topological properties or meet strict assumptions.

Work towards characterizing and understanding the identifiability of level-1 networks (Solís-Lemus and Ané 2016; Solis-Lemus et al. 2020; Gross et al. 2021; Allman et al. 2022) has led to many reticulate inference methods assuming an underlying level-1 network (Huber et al. 2010; Solís-Lemus and Ané 2016; Allman et al. 2019; LeMay et al. 2021). These methods have surged in popularity recently, largely due to their computational tractability and—for SNaQ (Solís-Lemus and Ané 2016) and NANUQ (Allman et al. 2019)— their ability to account for discordance both from gene flow and incomplete lineage sorting. However, the level-1 assumption was routinely violated for networks generated under the birth-death-hybridization process, particularly for highly reticulate networks. Fortunately, in practice, methods that make this assumption are often used to either infer few reticulate events (e.g., Morales-Briones et al. 2018; Karimi et al. 2020; Myers et al. 2022), or researchers assume that gene flow events are primarily isolated within each major clade (e.g., Esquerré et al. 2022)—with both cases being more likely to have level-1 networks. Further, recent identifiability results for level-2 networks (Van Iersel et al. 2009; van Iersel et al. 2020, 2022) may prove useful in extending the level-1 limitation of some methods. Nonetheless, it will be important to assess how robust these methods are to model violations and characterize possible biases when estimating networks that are not level-1.

## 5 Conclusions

In this study, we have shown that the specific assumptions about reticulation and sampling can have important effects on distribution of generated phylogenetic networks from the birth-death-hybridization process. There are many mechanisms that leave a specific reticulate pattern (see table in Box 1); both the biological process and the sampling framework affect how reticulate patterns (lineage generative, degenerative, neutral) manifest on phylogenetic networks. Methods that can directly investigate which type of reticulation occurs on phylogenetic networks (Hibbins and Hahn 2019; Flouri et al. 2020) will be vital as we infer patterns of gene flow. Further, system-specific knowledge—factors like life history (Wolf et al. 2001; Rumpho et al. 2008; Wendel et al. 2009; Schulte et al. 2012; Montanari et al. 2016; Meier et al. 2019), geographic distributions (López-Caamal et al. 2014; Ottenburghs et al. 2017; Dolinay et al. 2021), and genomic architecture (Wendel and Cronn 2003; Ottenburghs et al. 2017; Edelman et al. 2019)—should be incorporated to help inform how to model and interpret each type of reticulation. The specific reticulation types affect diversification and topology of phylogenetic networks, potentially violating assumptions of some methods or biasing diversification rate estimates if not taken into account. Overall, our work highlights the importance of thinking carefully and deliberately about how we model gene flow with the birth-death-hybridization process.

## 6 Additional

## 6.1 Acknowledgements

We would like to thank members of the Heath Lab for helpful comments and feedback on the manuscript. We would also like to thank Huw Ogilvie and two anonymous reviewers for helpful comments and observations. Lastly, we would like to thank Claudia Solis-Lemus and George Tiley for inviting us to be a part of the special issue.

## 6.2 Data Availability Statement

All scripts used to simulate data, perform analyses, and create figures are available at: https://github.com/jjustison/BDH_simulation. All simulated data and supplemental material are available on Zenodo at: https://doi.org/10.5281/zenodo.8371004

