## Supplement for "Exploring the distribution of phylogenetic networks generated under a birth-death-hybridization process"

### S1 Supplementary Material

We can now relax the assumption that speciation is greater than extinction $\lambda > \mu$. Allowing $\lambda \leq \mu$ makes $\sigma_n \geq 1$ for some $n$. However, since $\nu_+ > \nu_-$, the value of $\sigma_n$ is a monotonically decreasing function of $n$ that has the limit:

$$\lim_{n \to \infty} \sigma_n = \frac{\nu_-}{\nu_+}. \tag{1}$$

This implies that there is a value $m$ where:

$$\sigma_n \geq 1, \;\; n \leq m \tag{2}$$

and:

$$\sigma_n < 1, \;\; n > m. \tag{3}$$

We can consider $P_n$ to be the product of all $\sigma_n > 1$ terms that would appear in $S_n$ (*i.e.*, $\sigma_i$ from $n+1 \leq i \leq m$):

$$P_n = \prod_{i=n+1}^{m} \sigma_i. \tag{4}$$

Factoring $P_n$ out of the summation in $S_n$ results in summands that are now the product of only $\sigma_n$ terms that are less than 1:

$$S_n = \frac{1}{\lambda_n} + \frac{1}{\lambda_n} P_n \sum_{i=1}^{\infty} \left( \frac{\prod_{j=n}^{i} \sigma_{j+1}}{P_n} \right). \tag{5}$$

Direct comparison with the following convergent geometric series denotes that the sum in $S_n$ (and consequently $S_n$ itself) must also converge:

$$\sum_{i=1}^{\infty} \left( \frac{\prod_{j=n}^{i} \sigma_{j+1}}{P_n} \right) \leq \sum_{i=0}^{\infty} (\sigma'')^i, \tag{6}$$

Where $\sigma''$ is the supremum value of $\sigma_n$ when $n > m$:

$$\sigma'' = sup_{\{n>m\}} \sigma_n. \tag{7}$$

Lastly, we can observe that terms with $\sigma_n > 1$ only appear in $S_i$ when $1 \leq i \leq m$, giving:

$$R = \sum_{i=1}^{m} S_i + \sum_{j=m+1}^{\infty} S_j, \tag{8}$$

which must have $R < \infty$ due to the first term $\sum_{i=1}^{m} S_i$ being a finite sum and the second term $\sum_{j=m+1}^{\infty} S_j$ converging by applying the same argument as when $\sum_{i=1}^{\infty} S_j$ converged when $\lambda > \mu$ (Section 2.3.2). Intuitively, the process tends towards extinction when the population is less than $m$, however, there is a nonzero probability that the process will exceed $m$ where it will drift towards infinity in finite time.

#### S1.2 Supplementary Figures

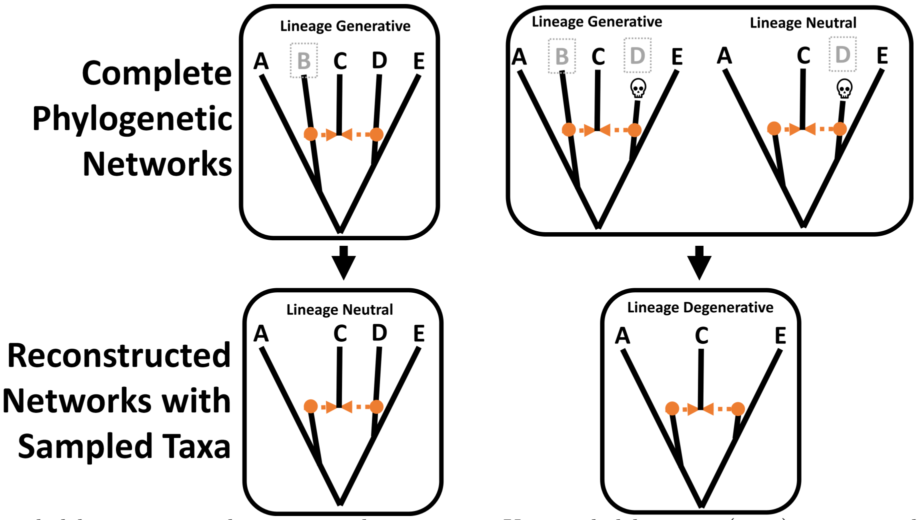


Figure S1: Unsampled lineages can change reticulation type. Unsampled lineages (grey) are pruned in the reconstructed phylogenies. Extinct species are depicted with a skull.

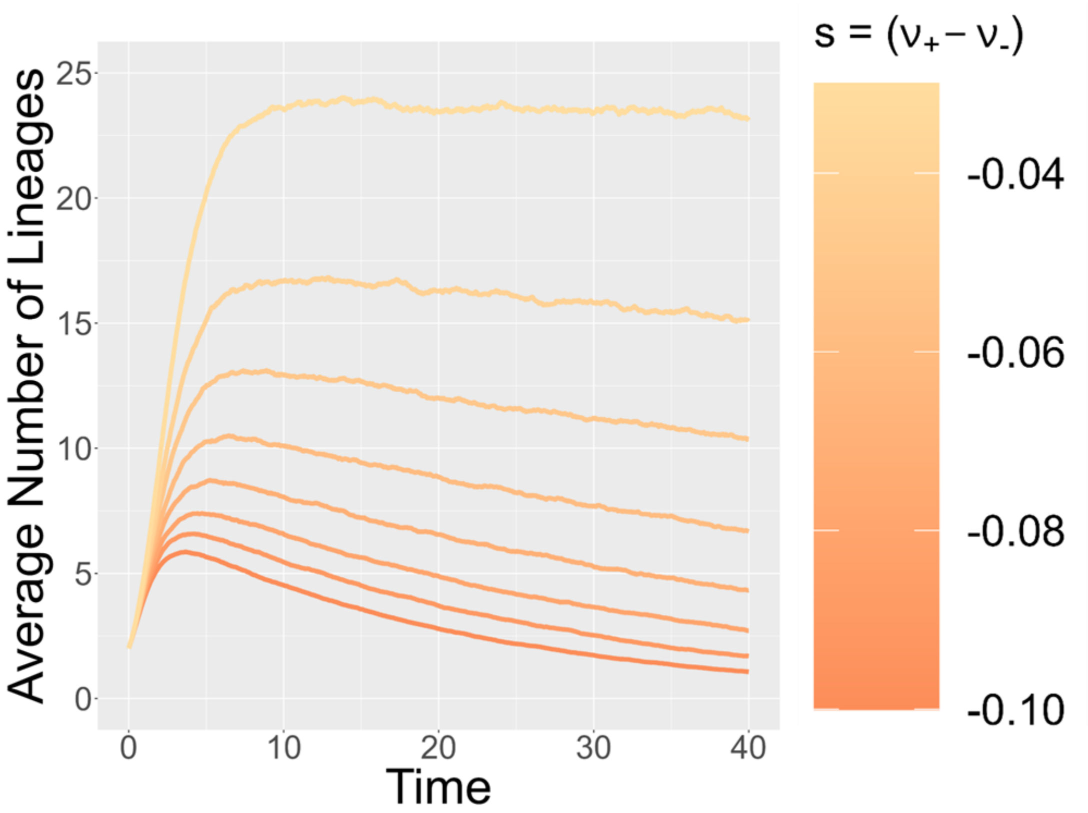


Figure S2: The average number of lineages through time with respect to $s$. We assume a speciation rate of $\lambda = 4$, extinction rate of $\mu = 3$, a hybridization rate of $\nu = 0.2$, and a lineage neutral hybridization rate of $\nu_0 = 0$. Higher lineage-pair diversification $s$ corresponds to a lighter orange, while low values of $s$ are denoted with a dark orange. Lines were only drawn for time points that had at least 4,500 replicates that had survived to that time point and had not reached the timeout time.

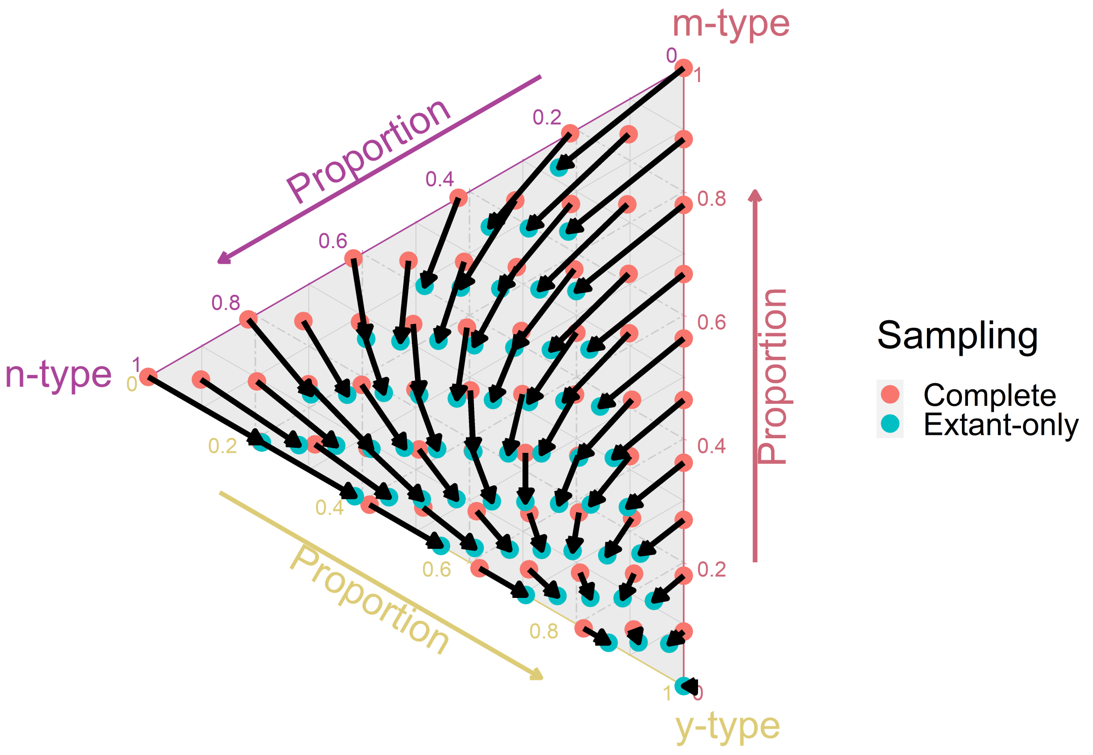


Figure S3: Observed hybridization types on simulated phylogenetic networks. Axes represent the observed proportion of total hybridizations for each hybridization type: lineage generative (m-type), lineage neutral (n-type), and lineage degenerative (y-type). Each circle represents the average proportion 20,000 replicates of a given hybrid rate combination. Red circles denote completely sampled phylogenies, while blue circles denote extant-only sampling. Circles connected with an arrow denote the exact same simulation conditions, with the exception of sampling.

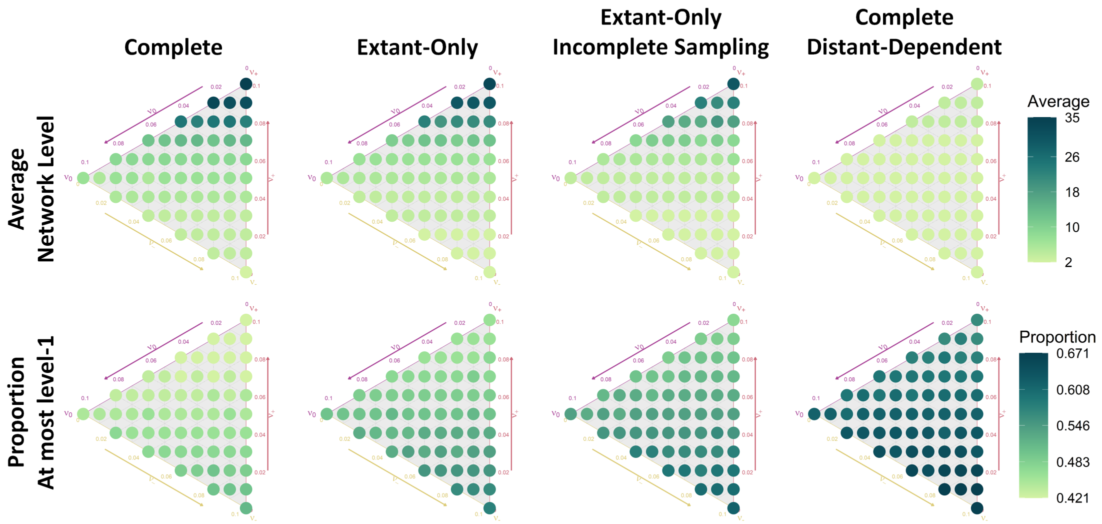


Figure S4: Phylogenetic level across the hybrid rate simplex. Each column denotes the simulation conditions used to simulate phylogenetic networks across the hybrid rate simplex (Section 2.4.2). Each row summarizes the average network level and proportion of phylogenetic networks that were level-1 or lower. The color of each point is used to depict the average value observed from 20,000, with darker colors representing a higher values.

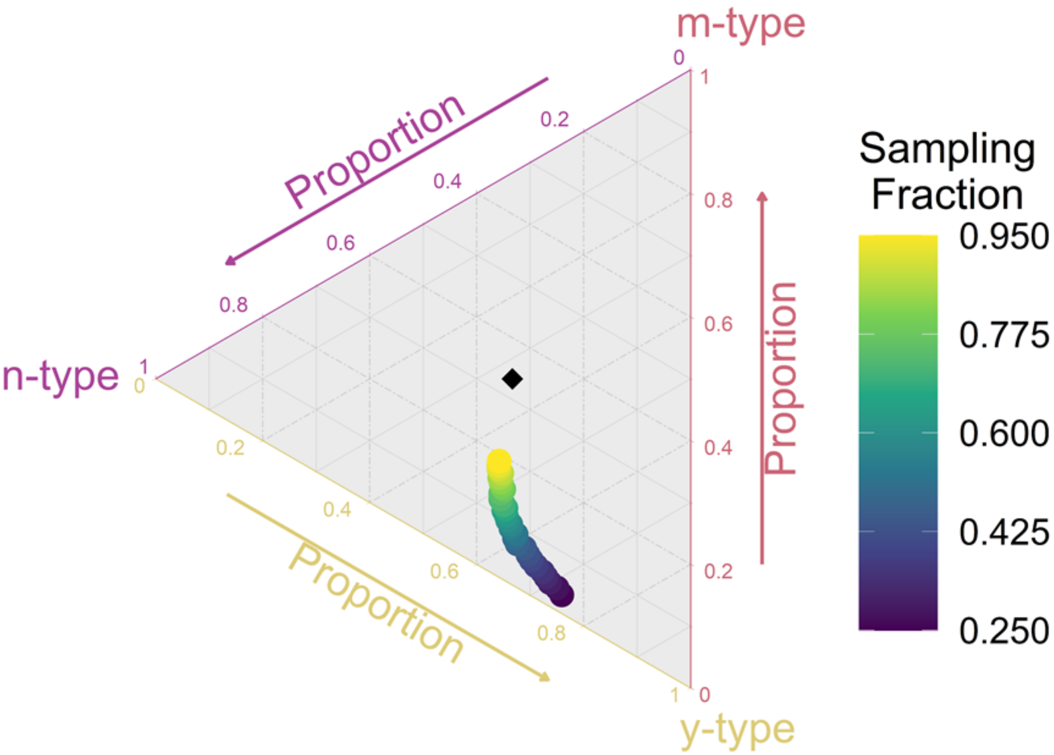


Figure S5: Observed hybridization types on simulated phylogenetic networks. Axes represent the observed proportion of total hybridizations for each hybridization type: lineage generative (m-type), lineage neutral (n-type), and lineage degenerative (y-type). Each circle represents the average proportion 20,000 replicates of a given hybrid rate combination. The black diamond represents the expected proportion with complete sampling.

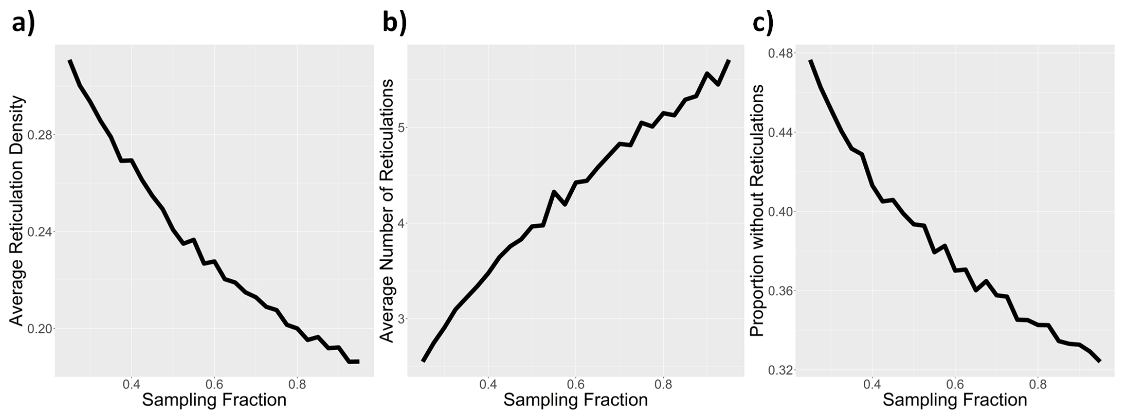


Figure S6: Reticulation summary statistics as a function of incomplete sampling. Values represent observed averages taken from 20,000 simulated phylogenetic networks for each sampling fraction value. The sampling fraction represents the number of extant tips that are randomly sampled from the network. Extant-only networks were used for the incomplete sampling dataset, while complete phylogenetic networks with extant and extinct species were used for the genetic distance-dependent dataset

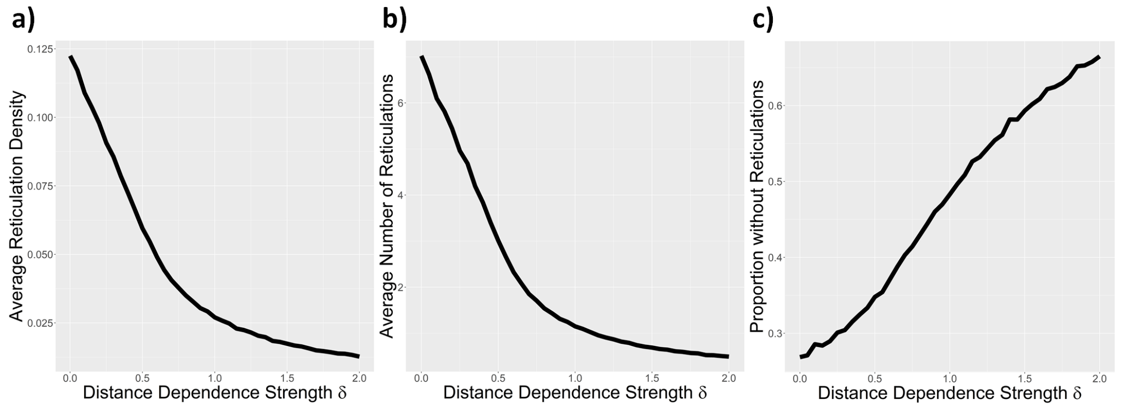


Figure S7: Reticulation summary statistics as a function of genetic distance-dependence. Values represent the observed averages taken from 20,000 simulated phylogenetic networks for each genetic distance-dependence strength ($\delta$). The value of *delta* corresponds to the strength of the genetic distance dependence, with higher values of $\delta$ meaning reticulation events primarily only occur between closely related lineages.

#### S1.3 Supplementary Tables

Table S1: Hybridization-type proportions. The overall hybridization rate ($\nu$) can be split into three separate rates: lineage generative ($\nu_+$), lineage neutral ($\nu_0$), and lineage degenerative ($\nu_-$). The values in the table represent the proportion of $\nu = 0.1$ for each hybridization type for simulations across the hybridization rate simplex.

| | $\nu_+$ | $\nu_-$ | $\nu_0$ |
|---|---|---|---|
| 1 | 0.00 | 1.00 | 0.00 |
| 2 | 0.10 | 0.90 | 0.00 |
| 3 | 0.05 | 0.85 | 0.10 |
| 4 | 0.00 | 0.80 | 0.20 |
| 5 | 0.20 | 0.80 | 0.00 |
| 6 | 0.15 | 0.75 | 0.10 |
| 7 | 0.10 | 0.70 | 0.20 |
| 8 | 0.05 | 0.65 | 0.30 |
| 9 | 0.00 | 0.60 | 0.40 |
| 10 | 0.30 | 0.70 | 0.00 |
| 11 | 0.25 | 0.65 | 0.10 |
| 12 | 0.20 | 0.60 | 0.20 |
| 13 | 0.15 | 0.55 | 0.30 |
| 14 | 0.10 | 0.50 | 0.40 |
| 15 | 0.05 | 0.45 | 0.50 |
| 16 | 0.00 | 0.40 | 0.60 |
| 17 | 0.40 | 0.60 | 0.00 |
| 18 | 0.35 | 0.55 | 0.10 |
| 19 | 0.30 | 0.50 | 0.20 |
| 20 | 0.25 | 0.45 | 0.30 |
| 21 | 0.20 | 0.40 | 0.40 |
| 22 | 0.15 | 0.35 | 0.50 |
| 23 | 0.10 | 0.30 | 0.60 |
| 24 | 0.05 | 0.25 | 0.70 |
| 25 | 0.00 | 0.20 | 0.80 |
| 26 | 0.50 | 0.50 | 0.00 |
| 27 | 0.45 | 0.45 | 0.10 |
| 28 | 0.40 | 0.40 | 0.20 |
| 29 | 0.35 | 0.35 | 0.30 |
| 30 | 0.30 | 0.30 | 0.40 |
| 31 | 0.25 | 0.25 | 0.50 |
| 32 | 0.20 | 0.20 | 0.60 |
| 33 | 0.15 | 0.15 | 0.70 |
| 34 | 0.10 | 0.10 | 0.80 |
| 35 | 0.05 | 0.05 | 0.90 |
| 36 | 0.00 | 0.00 | 1.00 |
| 37 | 0.60 | 0.40 | 0.00 |
| 38 | 0.55 | 0.35 | 0.10 |
| 39 | 0.50 | 0.30 | 0.20 |
| 40 | 0.45 | 0.25 | 0.30 |
| 41 | 0.40 | 0.20 | 0.40 |
| 42 | 0.35 | 0.15 | 0.50 |
| 43 | 0.30 | 0.10 | 0.60 |
| 44 | 0.25 | 0.05 | 0.70 |
| 45 | 0.20 | 0.00 | 0.80 |
| 46 | 0.70 | 0.30 | 0.00 |
| 47 | 0.65 | 0.25 | 0.10 |
| 48 | 0.60 | 0.20 | 0.20 |
| 49 | 0.55 | 0.15 | 0.30 |
| 50 | 0.50 | 0.10 | 0.40 |
| 51 | 0.45 | 0.05 | 0.50 |
| 52 | 0.40 | 0.00 | 0.60 |
| 53 | 0.80 | 0.20 | 0.00 |
| 54 | 0.75 | 0.15 | 0.10 |
| 55 | 0.70 | 0.10 | 0.20 |
| 56 | 0.65 | 0.05 | 0.30 |
| 57 | 0.60 | 0.00 | 0.40 |
| 58 | 0.90 | 0.10 | 0.00 |
| 59 | 0.85 | 0.05 | 0.10 |
| 60 | 0.80 | 0.00 | 0.20 |
| 61 | 1.00 | 0.00 | 0.00 |

Table S2: Parameters common to all simulations

| | |
|---|---|
| **Diversification Parameters** | $\lambda = 4$<br>$\mu = 2$<br>$\nu = 0.1$ |
| **Simulation Settings** | $age = 1$<br>20,000 replicates |
| **Rejection Conditions** | Extinct Phylogeny<br>Only one extant taxon<br>Simulation $>$ 10 seconds |

Table S3: Simulations across the hybrid-rate simplex. Each combination of hybridization rates is given in Table S1 and visually depicted in Fig. 3.

| Dataset | Parameters |
|---|---|
| **Complete** | **Sampling Settings**<br>Extinct species kept<br>$Sampling\ fraction = 1$<br><br>**Genetic Distance**<br>$\delta = 0$ |
| **Distance Dependent** | **Sampling Settings**<br>Extinct species kept<br>$Sampling\ fraction = 1$<br><br>**Genetic Distance**<br>$\delta = 0.5$ |
| **Extant-only** | **Sampling Settings**<br>Extinct species pruned<br>$Sampling\ fraction = 1$<br><br>**Genetic Distance**<br>$\delta = 0$ |
| **Incomplete Sampling** | **Sampling Settings**<br>Extinct species pruned<br>$Sampling\ fraction = 0.75$<br><br>**Genetic Distance**<br>$\delta = 0$ |

Table S4: Simulations using constant hybridization rates for each reticulation type ($\nu_+ = \nu_- = \nu_0 = \frac{1}{30}$).

| Dataset | Parameters |
|---|---|
| **Distance Dependent** | **Sampling Settings**<br>Extinct species kept<br>$Sampling\ fraction = 1$<br><br>**Genetic Distance**<br>$\delta$ varies from 0 to 2.0 |
| **Incomplete Sampling** | **Sampling Settings**<br>Extinct species pruned<br>$Sampling\ fraction$ from 0.20 to 1.0<br><br>**Genetic Distance**<br>$\delta = 0$ |

Table S5: Summary statistics across the hybrid simplex. Summary statistics were computed for each dataset by first taking the average value or proportion within each of the 61 hybrid rate proportions and then summarizing between each hybrid rate proportion (see Fig. 3 for hybrid rate proportions).

| | **Tips** | | | | **Reticulations** | | | | **Reticulation Density** | | | | **Proportion with Zero Reticulations** | | | |
|---|---|---|---|---|---|---|---|---|---|---|---|---|---|---|---|---|
| **Dataset** | mean | var | min | max | mean | var | min | max | mean | var | min | max | mean | var | min | max |
| **Complete** | 38.14 | 141.68 | 24.89 | 78.65 | 10.20 | 61.14 | 3.57 | 34.80 | 0.13 | 1.26e-4 | 0.11 | 0.15 | 0.27 | 0.00 | 0.26 | 0.28 |
| Extant-Only | 22.25 | 97.42 | 12.39 | 58.05 | 8.79 | 55.40 | 2.72 | 32.62 | 0.18 | 5.52e-05 | 0.17 | 0.20 | 0.32 | 0.00 | 0.31 | 0.34 |
| **Incomplete Sampling** | 16.51 | 48.26 | 9.41 | 43.82 | 7.36 | 35.50 | 2.34 | 29.13 | 0.21 | 8.84e-5 | 0.19 | 0.23 | 0.35 | 0.00 | 0.34 | 0.37 |
| **Distance Dependence** | 34.77 | 13.89 | 28.37 | 44.79 | 3.11 | 0.39 | 2.13 | 4.86 | 0.06 | 5.89e-7 | 0.06 | 0.06 | 0.35 | 0.00 | 0.34 | 0.36 |

Table S6: Class proportions across the hybrid simplex. Here, each rate combination is summarized and used as a point in analysis.

| | **Tree-based** | | | **FU-Stable** | | | **Tree-Child** | | | **Normal** | | |
|---|---|---|---|---|---|---|---|---|---|---|---|---|
| **Dataset** | mean | min | max | mean | min | max | mean | min | max | mean | min | max |
| **Complete** | 0.971 | 0.940 | 1.000 | 0.757 | 0.738 | 1.000 | 0.631 | 0.589 | 1.000 | 0.536 | 0.464 | 1.000 |
| **Extant-Only** | 0.959 | 0.931 | 0.996 | 0.737 | 0.722 | 0.810 | 0.601 | 0.582 | 0.702 | 0.486 | 0.452 | 0.613 |
| **Incomplete Sampling** | 0.951 | 0.922 | 0.967 | 0.733 | 0.709 | 0.773 | 0.598 | 0.588 | 0.624 | 0.483 | 0.468 | 0.535 |
| **Distance Dependence** | 0.997 | 0.995 | 1.000 | 0.903 | 0.882 | 1.000 | 0.789 | 0.710 | 1.000 | 0.658 | 0.556 | 1.000 |
